# Implant- and anesthesia-related factors affecting threshold intensities for vagus nerve stimulation

**DOI:** 10.1101/2021.01.22.427329

**Authors:** Umair Ahmed, Yao-Chuan Chang, Maria F. Lopez, Jason Wong, Timir Datta-Chaudhuri, Loren Rieth, Yousef Al-Abed, Stavros Zanos

**Author notes:** **Corresponding author** Stavros Zanos MD, PhD, Address: 350 community dr, Manhasset, New York, 11030, USA. **Authorship statement** UA conceived and designed experiments, performed experiments, analyzed and interpreted experimental results, and wrote the manuscript. YCC, MFL and JW performed experiments. TDC, LR and YAA critically reviewed and edited the manuscript. SZ conceived and designed experiments, analyzed and interpreted experimental results, wrote the manuscript, and obtained funding.

## Abstract

Vagus nerve stimulation (VNS) is used as therapy in epilepsy and depression and is tested as a potential treatment for several chronic disorders. Typically, VNS is delivered at increasing stimulus intensity until a response is observed (threshold intensity). Factors that affect threshold intensities for engagement of different fiber types and concomitant physiological responses have not been studied. We determined neural and physiological responses to increasing stimulus intensities of VNS in anesthetized and awake animals, and examined the effect of implant- and anesthesia-related factors on threshold intensities in a rodent model of VNS. In acute and long-term cervical vagus nerve implants (53 and 14 rats, respectively) VNS was delivered under isoflurane, ketamine-xylazine, or awake at different intensities. Stimulus-evoked compound action potentials (eCAPs) were recorded, elicited physiological responses were registered, including changes heart rate (HR), breathing, and blood pressure (BP), and threshold intensities were determined. The intensity that elicits eCAPs (“neural threshold”) is significantly lower than what elicits a physiological response (“physiological threshold”, PT) (25 μA ±1.8 vs. 70 μA ±5.2, respectively; Mean ±SEM). Changes in BP occur at the lowest stimulus intensities (80 μA ±7), followed by changes in HR (105 μA ±8.4) and finally in breathing (310 μA ±32.5). PT is lower with than without electrode insulation (60 μA ±12, vs. 700 μA ±123). PT and electrode impedance are correlated in long-term (*r*=0.47; *p*<0.001) but not in acute implants (*r*=-0.34; *p* NS); both PT and impedance increase with implant age (Pearson correlation *r*=0.44; *p*<0.001 and r=0.64; p<0.001, respectively). PT is lowest when animals are awake (210 μA ±33; Mean ±SEM), followed by ketamine-xylazine (630 μA ±154), and isoflurane (1075 μA ±131). The sequence of physiological responses with increasing VNS intensity is similar in both anesthetized and awake states. Implant age, electrical impedance and the type of anesthesia affect VNS threshold and should be accounted for when determining stimulation dose.

## Introduction

In neuromodulation, electrical energy is delivered with the intent of altering the function of the central or peripheral nervous system. It can be delivered either by means of invasive (e.g., deep brain stimulation and peripheral nerve stimulation)^1–4^ or non-invasive stimulation (e.g., transcranial stimulation and transauricular vagus nerve stimulation)^5–7^. Usually, neurostimulation is initiated at a low intensity, which is increased gradually until a neurophysiological, behavioral, or physiological response is observed. The minimum intensity to elicit such a response is defined as the stimulation threshold. At the cellular level, the stimulation threshold corresponds to the electrical energy level that is just enough to activate neural elements, typically neurons and their axonal and dendritic processes, with the lowest excitability threshold^8, 9^. At higher intensities, elements with higher excitability thresholds get recruited, leading to increasingly stronger or additional responses^8, 10^.

The type of threshold responses and the way they are measured depends on the site of neurostimulation. Examples of threshold responses are the motor evoked potential (MEP), a neurophysiological response to cortical stimulation^11^, reduction in pain, a behavioral response to spinal cord stimulation^12^, and a change in heart rate, a physiological response to cervical vagus nerve stimulation (VNS)^13^. These examples underscore that stimulation threshold responses depend on which components of the nervous system are activated or inhibited, but also on the end-organs through which that activation, or inhibition, manifests itself. For example, motor cortex stimulation would not elicit the same MEP in subjects taking muscle relaxants and in subjects not taking any medication^14^. Similarly, the behavioral response to spinal cord stimulation depends on cognitive state^15^. Therefore, the physiological state of the neural circuits and end-organs is important when the stimulation threshold is determined.

Knowledge of threshold intensities is essential in neuromodulation studies and therapies. The threshold for a given stimulation modality varies widely across subjects^16, 17^. Mechanisms that determine threshold can be grouped into 3 categories: those related to the neural interface, those related to the generated electrical field, and those related to the physiological status of the end-organ. Factors that determine the neural interface include surgical technique^2, 11^, foreign body response to a long-term implanted electrode^1, 8 19^, and degradation of the electrode itself^20^. Factors influencing the generated electrical field include electrode geometry^21 22^, electrode orientation relative to the stimulated neural elements^17, 21, 23^, and stimulation parameters^11, 21^. Finally, the state of a subject or the end-organ at the time of stimulation affects the stimulation threshold^24^. For example, general anesthesia has distinct effects on the nervous system and multiple end-organs, altering stimulation thresholds through several mechanisms^24^. Factors affecting stimulation thresholds in brain stimulation^2, 11 21^, spinal cord stimulation^23^, and somatic peripheral nerve stimulation, such as tibial or ulnar nerve^18, 20, 24^ are relatively well-studied. There is much less data on factors affecting stimulation thresholds in vagus nerve stimulation (VNS), an established neuromodulation therapy in epilepsy and depression^1, 25, 26^, and currently under study in a variety of cardiovascular, immune, and metabolic disorders^27–30^.

In this study, we examined several factors affecting threshold intensity in cervical VNS in rats, and more specifically the presence of insulation at the neural interface, the level of *ex-vivo* and *in-vivo* electrode impedance, the age of the implant, and the conditions of anesthesia. To address these questions, we developed acute and long-term implants of vagus nerve stimulation and recording, and a methodology to determine thresholds in individual animals. We found that the threshold for evoking nerve compound action potentials with VNS (“neural threshold”, NT) is lower than that for eliciting a reproducible physiological response (“physiological threshold”, PT). The PT has a weak correlation with *ex-vivo* or *in-vivo* electrode impedance on an individual measurement basis, even though they both increased with implant age. Anesthesia caused a significant increase in the threshold of VNS, compared to the conscious state: isoflurane has the largest effect, followed by ketamine-xylazine.

## Methods

### Animal preparation

Male Sprague-Dawley rats (n=67) weighing 250-400gm were used in this study. Institutional Animal Care and Use Committee approved all animal experiments. lsoflurane was a primary anesthetic agent (induction 4%, maintenance 1-2%). Ketamine (80 mg/kg IP) and xylazine (6 mg/kg IP) were used in a few experiments. After induction, rat was placed in a supine position, and hairs were removed from surgical sites. Alcohol and Betadine were used to disinfect surgical sites. Adequate level of anesthesia was assessed by pedal reflex and breathing rate.

### Instrumentation of physiological sensors

Physiological signals were monitored during experiments. Nasal temperature sensor was inserted inside nostril to detect temperature changes during inhalation and exhalation; breathing rate was calculated using nasal temperature sensor (fig. 1). Electrocardiogram (ECG) was recorded using 3-needle electrodes inserted intradermally (fig. 1). Left femoral artery was isolated, and polyethylene-tube 50 was inserted to monitor systemic arterial pressure (SAP) (fig. 1). Body temperature was measure using rectal probe and maintained at 37-37.5°c using heating pad.

**Figure 1.**
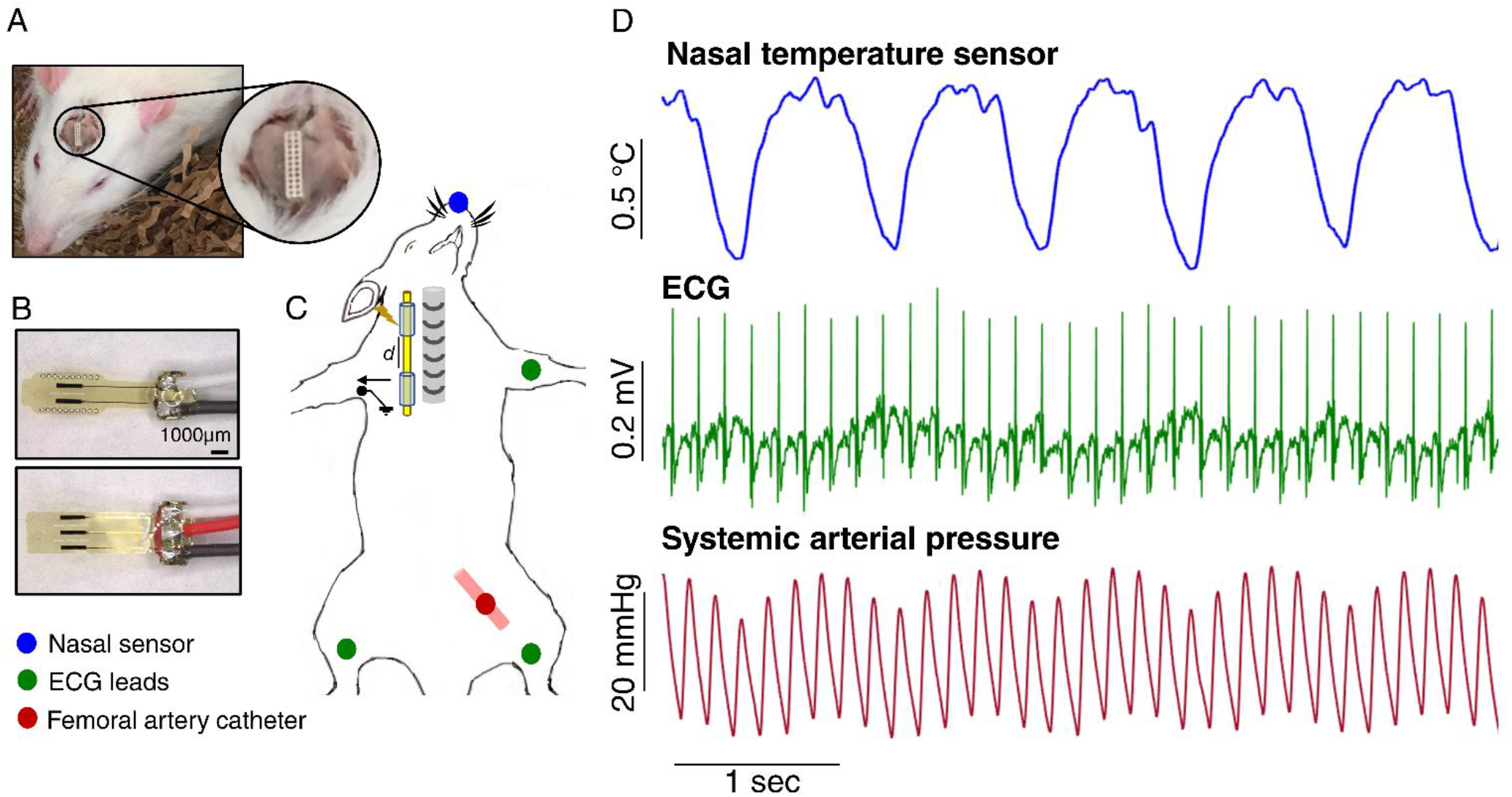
Schematic of an experiment in a rat, placement of electrodes, and instrumentation of physiological sensors. (A) Implanted head connector on the skull of a rat, VNS was delivered through this connector in a long-term vagal cuff implanted rat. (B) Close-up images of bipolar and tripolar Flex electrodes. (C) Two cuff electrodes were placed around a cervical vagus nerve (beside Trachea), one electrode was used for stimulation and the other to record stimulus-elicited evoked compound action potential (eCAP). Three needle Electrocardiogram (ECG) leads were placed intradermally on the limbs (green color), nasal temperature sensor was placed inside the nostril (blue color) and a catheter was inserted inside a femoral artery to measure systemic arterial pressure (red color). (D) Real time physiological signals acquired using the physiological sensors. Nasal airflow signal obtained from nasal temperature sensor (top panel), ECG signal from ECG leads (middle panel), and systemic arterial pressure from femoral artery catheter (bottom panel).

### Electrodes

The “Flex” electrodes used for this study were designed for acute and long-term implantation and made using microfabrication processes^31^ (fig. 1B). Some of the acute experiments were conducted using commercially available CorTec bipolar electrodes (CorTec, Germany). Additional microfabrication details of electrodes are described in supplementary file.

### Surgical procedure for acute implantation

A midline skin incision of 2-3 cm was given on the neck. After the incision, a submandibular gland can be visualized (fig. 2A). This gland was separated to visualize sternohyoid (midline) and sternocleidomastoid (oblique) muscles (fig. 2B, 2C). The sternocleidomastoid muscle was then retracted using magnetic retractors (Fine Science Tools, CA) to reach the carotid bundle (common carotid artery and vagus nerve; fig. 2C). A third muscle named omohyoid can be seen connecting sternohyoid and sternocleidomastoid muscles; carotid bundle can be visualized rostral and caudal to the omohyoid muscle (fig. 2C, 2D). A dissecting microscope (Carl Zeiss Microscopy, NY) was used to separate vagus nerve from carotid bundle (fig. 2D). The rostral end of vagus nerve was isolated, and in a few experiments, the caudal end was also isolated. The middle part of the nerve embedded under omohyoid muscle was left intact to minimize nerve manipulation. A bipolar or tripolar electrode was placed under the nerve (fig. 2E). In-case of two electrodes, the distance between them was always between 5-6 mm. A silicone elastomer Kwiksil (World Precision Instruments, FL) was used to insulate the nerve-electrode interface from the surrounding fluid.

**Figure 2.**
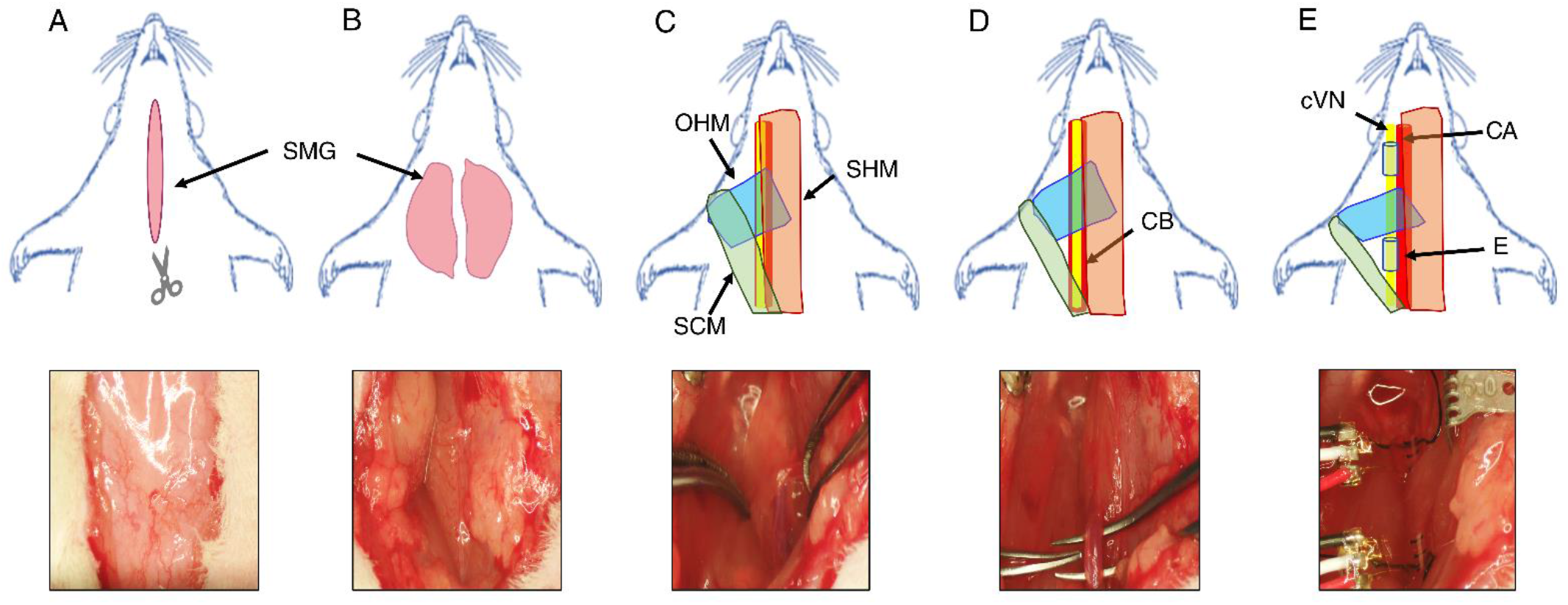
Surgical steps of a cervical vagus nerve isolation and electrode placement in a rat. (A) Midline neck incision while the rat was in a supine position. (B) Separation of submandibular salivary glands (SMG). (C) Major cervical muscles involved when isolating cervical vagus nerve (SCM = Sternocleidomastoid muscle; SHM = Sternohyoid muscle; OHM= Omohyoid muscle). (D) Retraction of SCM to better visualize carotid bundle (CB). (E) Placement of two cuff electrodes above and below OHM (cVN = Cervical vagus nerve; CA= Carotid artery; E = Electrode).

### Surgical procedure for long-term implantation

A 1 cm skin incision was given on the skull while the rat was in a prone position. Scalp tissue was cleared using hydrogen peroxide between bregma and lambda. A drilling machine using a bit of shaft diameter 2.3 mm (19007-09, FST, CA) was used to drill two holes on the right and left side of the skull, between coronal and lambdoid sutures. Self-tapping bone screws (19010-10, FST, CA) were screwed on these holes using a screwdriver (Stanley, CT). Rat was then placed in a left lateral recumbent position. A subcutaneous tunnel was then made between skull and neck using a hemostat, through which electrode leads were tunneled. Rat was then placed in a supine position, and electrode was placed around the left vagus nerve, as explained above. A small ligating clip (LT-100, Johnson & Johnson, NJ) was used to close Flex electrode. Electrode leads were sutured on a surrounding muscle to minimize nerve manipulation during physical movement. Skin was closed using 4.0 nylon-suture. Rat was then again placed in a prone position to fix the head-connector on the skull. A layer of acrylic dental cement (A-M systems, WA) was applied on the exposed area of skull, head connector was then fixed between the two skull screws (fig. 1A). Subcutaneous antibiotics (Enfloroxacin 5 mg/kg) and Meloxicam 3 mg/kg were given once a day for three days post-surgery. Few rats were purchased pre-implanted with a telemetry device (HD-S21, Data Science International, MN) to record the effect of VNS on systemic arterial pressure and heart rate on freely moving rats.

### Stimulation and Recording

All biological signals were first pre-processed using Bio Amplifier for ECG signals, Bridge Amplifier for pressure signals, and T-type Pods for nasal temperature sensor and rectal probe. These amplifiers and pods were linked to Powerlab (ADlnstruments Inc, CO). All the raw signals captured by Powerlab were displayed on LabChart (ADI, CO). The biological signals from telemetry implanted rats were captured on PhysioTel Hardware (OSI, MN), and were displayed on LabChart. The stimulator (STG4008, Multi-Channel Systems, DE) was connected to the rostral electrode, and a recording amplifier (RHS2000 series, lntan Technologies, CA) was connected to the caudal electrode. Physiological threshold (PT) was defined as intensity needed to elicit a 5-10% decrease in heart rate (HR). Neural threshold (NT) was defined as intensity needed to elicit a visible eCAP on a recording electrode. Stimulation parameters used for determining thresholds were constant current, rectangular pulses with pulse widths (PW) of 100 μs, frequency (F) of 30 Hz, and duration of 10 s. Stimulus triggers were registered on LabChart as well as on lntan software and were used to synchronize the recordings. Parameters used in this study are PW of 100 μs, F= 30 Hz, and duration 10 s. Bipolar stimulation was used in terminal and survival experiments.

### Analysis

#### Analysis of physiological and neurophysiological signals

Physiological and neural recording data were processed using custom scripts on MATLAB (Mathworks). The ECG signal was first pre-processed with a high-pass filter at 0.1 Hz to remove direct current (DC) voltage drift, followed by QRS peak detection to calculate heart rate (HR) based on individual R-R intervals. For nasal temperature sensor signal, a high-pass filter at 0.1 Hz was applied to eliminate the temperature drift. Both positive and negative peaks were detected to compute breathing cycles for accurate detection of each breath.

The change in breathing rate (ΔBR) and heart rate (ΔHR) for stimulation trains were calculated as follow:

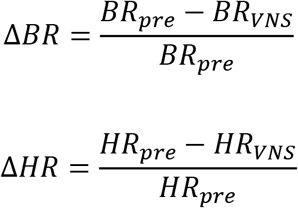

where BR_pre_ (or HR_pre_) corresponds to the mean BR (or HR) during the 1Os long train before the stimulation, and BR_VNS_ (or HR_VNS_) to the mean BR (or HR) during the stimulation. Neural signal processing and elimination of non-neural signals captured from a recording nerve electrode. Briefly, non-neural components in eCAP recordings include ECG-artifact, stimulus-artifact, and EMG-artifact from laryngeal muscle contraction. To improve nerve signal quality and resolve short-latency fiber responses, a stimulus artifact suppression algorithm was employed.

#### Statistical analysis

A paired *t*-test was used to compare the physiological and neural thresholds. A paired *t*-test was used to compare physiological thresholds of animals under isoflurane, ketamine-xylazine, and awake state. Pearson-correlation coefficient was used to find correlation between physiological and neural thresholds, and between physiological threshold and electrode impedance. Comparisons were deemed statistically significant for *p*<0.05 for all analyses. All statistical analyses were conducted on MATLAB (Mathworks).

## Results

### Neural threshold is significantly lower than physiological threshold

We sought to investigate the relationship between the minimum stimulus intensity that elicits a neural response (“neural threshold”, NT) and the minimum intensity that elicits a vagally-mediated physiological response (“physiological threshold”, PT). We determined NT by measuring stimulus-evoked compound action potentials (eCAPs) (fig. 3A). We determined PT by measuring stimulus-elicited change in heart rate (HR). eCAP responses comprise components with different latencies, corresponding to activation of different fiber types, recruited at different intensities: short latency A-fibers are engaged at low intensities, whereas intermediate latency B-fibers are engaged at higher intensities (fig. 3A). This fiber recruitment pattern correlates well with the physiological responses seen during VNS: at low-intensities, A-fibers mediate an increase in SAP and moderate drop in BR, at intermediate-intensities B-fibers mediate drop in HR, and secondarily in SAP, and at high intensities C-fibers mediate apnea responses (fig. 3A, 4).

**Figure 3.**
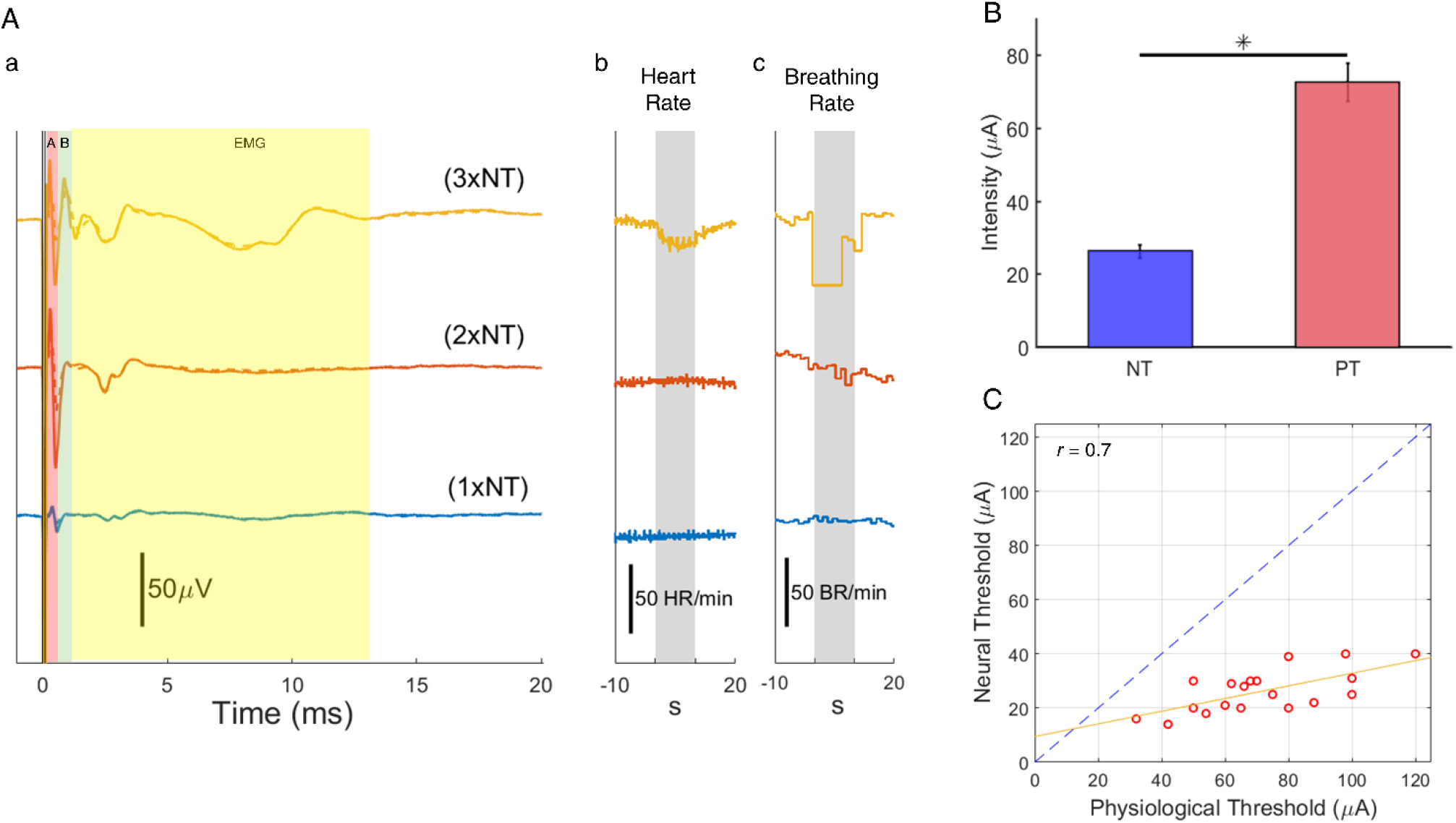
Relation between neural and physiological thresholds of VNS. (A) Examples of evoked compound action potentials (eCAP) (a) and stimulus-elicited changes in heart rate (b) and breathing rate (c), at different stimulus intensities, expressed in units of neural threshold (1xNT, 2xNT and 3xNT). To determine NT, trains of 10 s duration, 30 Hz, of 100 ms pulse width were delivered. The colored shaded areas in panel ‘a’ represent the latency windows used for quantifying magnitude of fiber activation, based on fiber conduction velocity and electrode distance; in this case, the window for A-fibers was 0.3 - 0.95 ms, for B-fibers 0.7 - 1.2 ms, and for evoked EMG activity (due to efferent, laryngeal A-fiber activation) 2 - 13 ms. The grey shaded area in panels ‘b’ and ‘c’ represent the duration of stimulation. (B) Mean (±SEM) neural and physiological thresholds in 19 animals (asterisk indicates *p*<0.001, Paired *t*-test). (C) Comparison between NT and PT values in individual animals (n=19). Each point in the plot represents an experiment in which both NT and PT were determined. Dotted blue line represents unity line, yellow line represents the linear fit of the data. NT and PT are significantly correlated (Pearson correlation coefficient: *r*=0.7, *p*<0.001).

**Figure 4.**
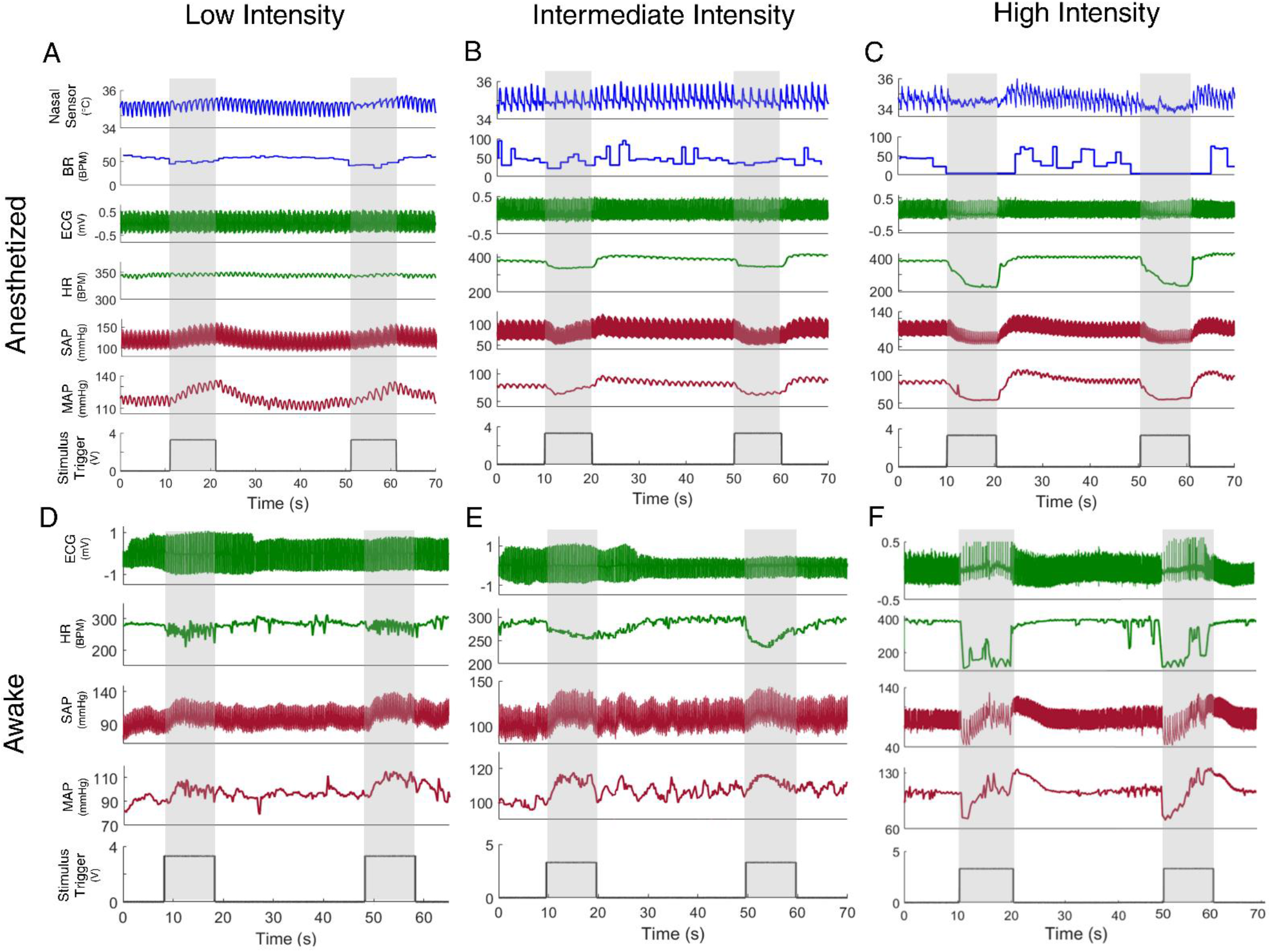
Examples of cardiopulmonary responses to VNS of different intensities, in anesthetized and awake rats. (A) Cardiopulmonary responses of VNS registered in a rat, anesthetized with isoflurane. From top to bottom panel: nasal airflow, breathing rate (BR), ECG, heart rate (HR), systemic arterial pressure (SAP), mean arterial pressure (MAP), and stimulus trigger. Two 1a-seconds stimulus trains were delivered with similar parameters (grey shaded area). Low intensity stimulation parameters are 80μA, 30Hz, 100μs. (B) Same as (A), except intermediate stimulation intensity (280μA, 30Hz, 100μs). (C) Same as (A), except high stimulation intensity (700μA, 30Hz, 100μs). (D) Physiological responses of VNS in awake, freely-moving rat. From top to bottom panel: ECG, HR, SAP, MAP, and stimulation trigger. Two 1a-seconds stimulus trains were delivered with similar parameters. Low intensity stimulation parameters are 50μA, 30Hz, 100μs. (E) Same as (D), except intermediate stimulation intensity (250μA, 30 Hz, 100μs). (F) Same as (D), except high stimulation intensity (600μA, 30Hz, 100μs).

In 19 animals, NT range between 14 and 40 μA and PT between 42 and 120 μA. Mean NT (25 μA± 1.83) is significantly smaller than mean PT (70 μA± 5.24) (p<0.01, paired *t*-test) (fig. 3B). On a single animal basis, NT is always lower than PT (fig. 3C), and NT and PT were correlated: animals with relatively low NT values have relatively low PT values, and vice-versa (Pearson-correlation coefficient: *r*=0.7, p<0.01). NT and PT are not significantly different between right and left VNS (Suppl. fig. S1). In a separate group of 4 rats, neural and physiological responses were compared by using 2 different stimulation frequencies, 1 and 30 Hz; neural responses are similar in both frequencies, whereas physiological responses are significantly different (table I). These results indicate that there is a significant, almost 3-fold, variability in both NT and PT across animals and that PT is 2-4 times greater than NT. These results also suggest that stimulation frequency of VNS affects the magnitude of the physiological response, but not of the neural response.

**Table I.** Effect of stimulation frequencies on neural and physiological responses. Mean (±SEM) amplitude of A- and B-fiber response, extracted from eCAPS, and magnitude of stimulus-elicited changes in heart rate (ΔHR), for 2 different stimulation frequencies, 1 and 30 Hz. Stimulation intensities and pulse widths are identical for both frequencies (n=4). Asterisks denote statistically significant difference between the 2 frequencies (*p*<0.001, Paired *t*-test).

| Frequency<br>(Hz) | A-fiber<br>($\mu\text{V}$) | B-fiber<br>($\mu\text{V}$) | $\Delta\text{HR}$<br>(%) |
| --- | --- | --- | --- |
| 1 | 121.42 ± 11.2 | 40.87 ± 9.6 | -0.08 ± 0.2* |
| 30 | 138.27 ± 11.5 | 37.55 ± 8.6 | -17.30 ± 3.3* |

### Acute cardiopulmonary responses to VNS depend on stimulus intensity

Different acute cardiopulmonary responses are elicited at different VNS intensities, including changes in breathing rate, usually in the form of slower breathing, and in heart rate, usually in the form of bradycardia (fig. 3A). Generally, intensity is a major determinant of the pattern of physiological responses to VNS. In animals anesthetized with isoflurane, at low VNS intensities, we routinely observed slower breathing and increase SAP, both of which returned to baseline after the end of stimulation (fig. 4A). At intermediate intensities, slower breathing and bradycardia are typically seen; often, the bradycardic response is associated with a drop in SAP; again, parameters returned to baseline after the end of stimulation (fig. 4B). At higher intensities, we observed apnea during stimulation, more intense bradycardia, and a larger drop in SAP; even though breathing and heart rhythm returned to baseline after the end of stimulation, a rebound increased in SAP is commonly seen (fig. 4C). Thresholds for different responses under anesthesia are summarized in table. II. Heart rate responses to VNS in awake, freely behaving animals are comparable to those in anesthetized animals, but SAP responses are often more complex and variable (fig. 4D-F). Overall trends for low, intermediate, and high intensities of anesthetized and awake groups are summarized in table. III. These results indicate that acute cardiopulmonary responses to VNS depend on the stimulus intensity.

**Table II.** Stimulation threshold intensities of various cardiopulmonary responses under anesthesia (isoflurane) in acute experiments (Mean ±SEM). Stimulation frequency and pulse width are 30 Hz and 100 μs, respectively (n=8 animals).

| **Mean Arterial Pressure**<br>**Threshold**<br>**($\mu\text{A}$)** | **Bradypnea**<br>**Threshold**<br>**($\mu\text{A}$)** | **Heart Rate**<br>**Threshold**<br>**($\mu\text{A}$)** | **Apnea**<br>**Threshold**<br>**($\mu\text{A}$)** |
| --- | --- | --- | --- |
| 80 ±7 | 80 ±7 | 105 ±8.4 | 310 ±32.5 |

**Table III.** Effect of stimulation intensity on cardiopulmonary responses in anesthetized (isoflurane) and in awake animals (Mean ±SEM). Stimulation frequency and pulse width are 30 Hz and 100 μs, respectively (n=4 animals for each group, anesthetized (isoflurane) and awake). PT=Physiological threshold.

|  | **Relative**<br>**Intensity**<br>**(xPT)** | **Absolute**<br>**intensity**<br>**(μA)** | **ΔBR**<br>**(%)** | **ΔHR**<br>**(%)** | **ΔMAP**<br>**(%)** |
| --- | --- | --- | --- | --- | --- |
| **Anesthetized** | Low<br>(0.5-1PT) | 80-100 | -13.84 ±8.1 | -0.74 ±1.1 | 10.09 ±2.8 |
| | Intermediate<br>(1-3PT) | 100-300 | -61.49 ±4.4 | -14.76 ±1.5 | -16.09 ±1.3 |
| | High<br>(>3PT) | 300-600 | -75.47 ±5.2 | -49.75 ±7.6 | -33.77 ±4.2 |
| **Awake** | Low<br>(0.5-1PT) | 20-70 | - | 0.26 ±0.7 | 16.85 ±3.6 |
| | Intermediate<br>(1-3PT) | 70-200 | - | -14.97 ±1.5 | 26.45 ±4.8 |
| | High<br>(>3PT) | 200-300 | - | -66.37 ±3.6 | -20.11 ±2.4 |

### Physiological threshold is lower with electrode insulation

In many acute experiments, we observed an increase in PT when moderate amounts of fluid accumulated around the stimulation electrode. We compared PT between experiments in the same animals with and without application of a silicone elastomer for additional insulation (Suppl. fig. S2). In the same 7 animals, PT with application of insulating elastomer is significantly lower than without: 60 μA ±12.3, vs. 700 μA ±122.6 (Suppl. table I). In one animal, we were able to reduce PT by dabbing the fluid from the area around the stimulating electrode with an absorbing cotton tip (Suppl. table I). These results indicate that additional insulation of the stimulating electrode prevents current shunting by accumulated fluid, resulting in lower PT values.

### Physiological threshold increases with increasing implant age

To determine whether implant age affects physiological responses to VNS, we chronically implanted stimulating electrodes on the cervical vagus and determined PT under anesthesia (isoflurane) in multiple stimulation sessions across several weeks until no physiological response was observed for two consecutive stimulation sessions. In general, the intensity to elicit the same physiological response is increased with increasing age; for example, to elicit a drop in HR by 8-10% 1 day after implantation intensity required is 100 μA, whereas at 60 days later, it is 1000 μA (fig. 5A, 5B). In 14 rats, PT is gradually increased significantly during the first 2 weeks, and only moderately afterward (fig. 5C). Mean PT at the day of implantation is 190 μA± 28.5 (n=14), 650 μA ±112.8 (n=12) at 2 weeks post-implantation, 670 μA ±100 (n=9) at 4 weeks, 630 μA ±160.7 (n=4) at 8 weeks, and 950 μA ± 550 (n=2) at 12 weeks. Additionally, significant amount of fibrosis is observed around the neural interface during the autopsy; the fibrotic response seems to get worse with the age of the implant (Suppl. fig. S3). These results indicate that implant age is positively correlated with PT, with the greatest increases happening during the first two weeks post-implantation.

**Figure 5.**
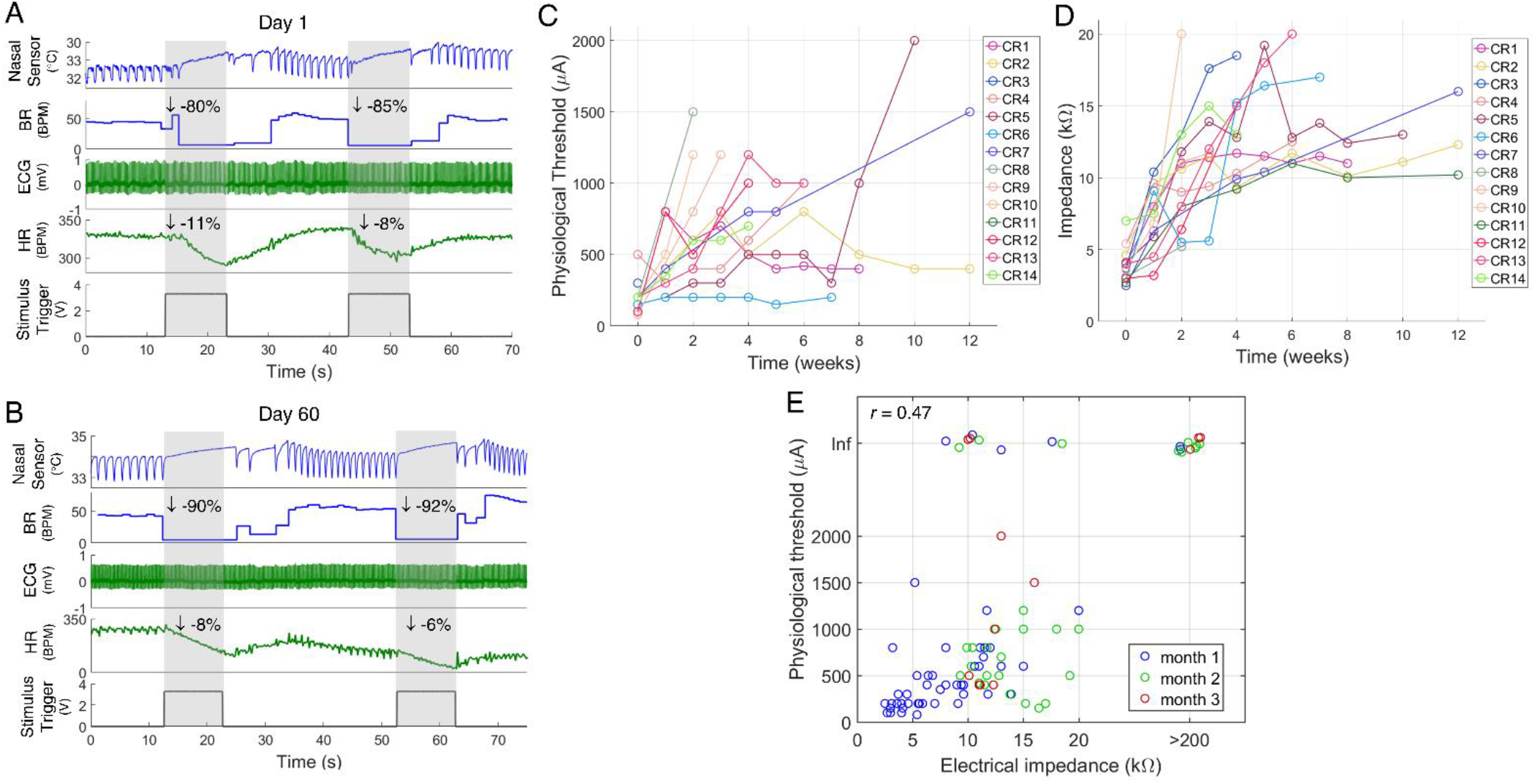
Effect of implant age on the physiological threshold and electrode impedance. (A) Example of a stimulus elicited change in heart rate and breathing rate at day 1 post-implantation under anesthesia (isoflurane). The grey shaded areas represent the duration of stimulation. Intensity used to elicit change in heart rate (ΔHR) and breathing rate (ΔBR) was 100μA (30Hz, 100μs PW). (B) Same animal as (A) but on day 60 of post-implantation. Intensity used to elicit ΔHR and ΔBR was 1000μA (30Hz, 100μs PW). (C) Longitudinal changes in physiological thresholds after implantation. Each trace represents one animal; the time point at which the line terminates corresponds to the loss of physiological response. (D) Longitudinal changes in electrode impedance after implantation. Each trace represents one animal; the time point at which the line terminates corresponds to impedance value >20 kΩ. (E) Impedance values vs. corresponding physiological threshold values from 14 animals. Each circle represents one experiment; different colors represent measurements performed at different implant ages: within 1 month from implantation (blue), 1-2 months (green), 2-3 months (red) Pearson correlation coefficient: *r* = 0.47; *p*<0.001). Inf denotes undeterminable PT (non-functional implant).

### Physiological threshold is affected by *in-vivo/ex-vivo* electrical impedance

To determine whether the level of the electrical impedance of the stimulating electrode, often used to infer the functional status of an electrode or the tissue-electrode interface, affects PT, we measured *in-vivo/ex-vivo* electrical impedance and PT in the same stimulation sessions, in acute and long-term nerve implants. In 53 acute experiments, mean *ex-vivo* impedance at 1KHz before implantation is 1.6 kΩ ±0.14, and mean PT is 80 μA ±5.84. The correlation between ex-vivo impedance and PT is non-significant (*r* = 0.21, *p* NS) (Suppl. fig. S4). In 12 animals, we compared the *ex-vivo* and *in-vivo* impedances of electrodes, measured immediately before and after implantation, respectively; *ex-vivo* impedance is always lower than the *in-vivo* impedance, and the two impedance values are highly correlated (*r* = 0.83) (Suppl. fig. S4). In 14 rats implanted with long-term electrodes, PT progressively increases with implant age (fig. 5C; *r* = 0.44, p<0.001). Electrode impedance also increases with age (fig. 5D) (*r* = 0.64, *p*<0.001). On a single measurement basis, there is a weak but statistically significant correlation between PT and impedance (fig. 5E; *r* = 0.47, *p*<0.001). In 3 out of 14 long-term implants, impedances were low but no physiological response was observed (fig 5E). These results indicate that *in-vivo/ex-vivo* electrical impedance of the stimulating electrode is a poor predictor of PT in acute implants. *In-vivo* impedance is a predictor in most, but not all, long-term implants.

### Physiological threshold is higher under anesthesia compared to wakefulness

To determine the effects of anesthesia on PT in rats chronically implanted with vagus electrodes, we measured PT under anesthesia with isoflurane, with ketamine-xylazine, and during awake conditions. In these experiments, the animal was first anesthetized with isoflurane, PT was measured, and isoflurane was turned off; after the animal recovered from isoflurane, PT was determined again, either in an awake state (first animal group; fig. 6A, 6B) or after a second round of anesthesia with ketamine-xylazine (second animal group). In the first group of 5 animals (23 experiments), PT measured under anesthesia with isoflurane is 1020 μA ±122.3, and in awake conditions, it is 210 μA ±33.5 (*p*<0.001, Paired *t*-test) (fig. 6C). In the second group of 5 animals (7 experiments), PT measured under isoflurane is 1215 μA ±207.5, and under ketamine-xylazine is 630 μA ±153.8 (*p*<0.001, Paired *t*-test) (fig. 6C). To determine whether these differences persisted at supra-threshold intensities, we measured the stimulus-elicited change in HR (ΔHR) to the same suprathreshold intensities, under isoflurane vs. awake, or under isoflurane vs. ketamine-xylazine. In the first group, the mean change in heart rate (ΔHR) under isoflurane is −2 % ±0.7, and in awake conditions, it is −26 % ±3.3 (*p*<0.01, paired *t*-test) (fig 6D). In the second group, HR under isoflurane is −1 % ±1.01 and under ketamine-xylazine it is −13 % ±3.98 (p<0.01, paired *t*-test) (fig. 6E). These results indicate that PT are the smallest when the animals are awake, followed by anesthesia with ketamine-xylazine and finally with isoflurane.

**Figure 6.**
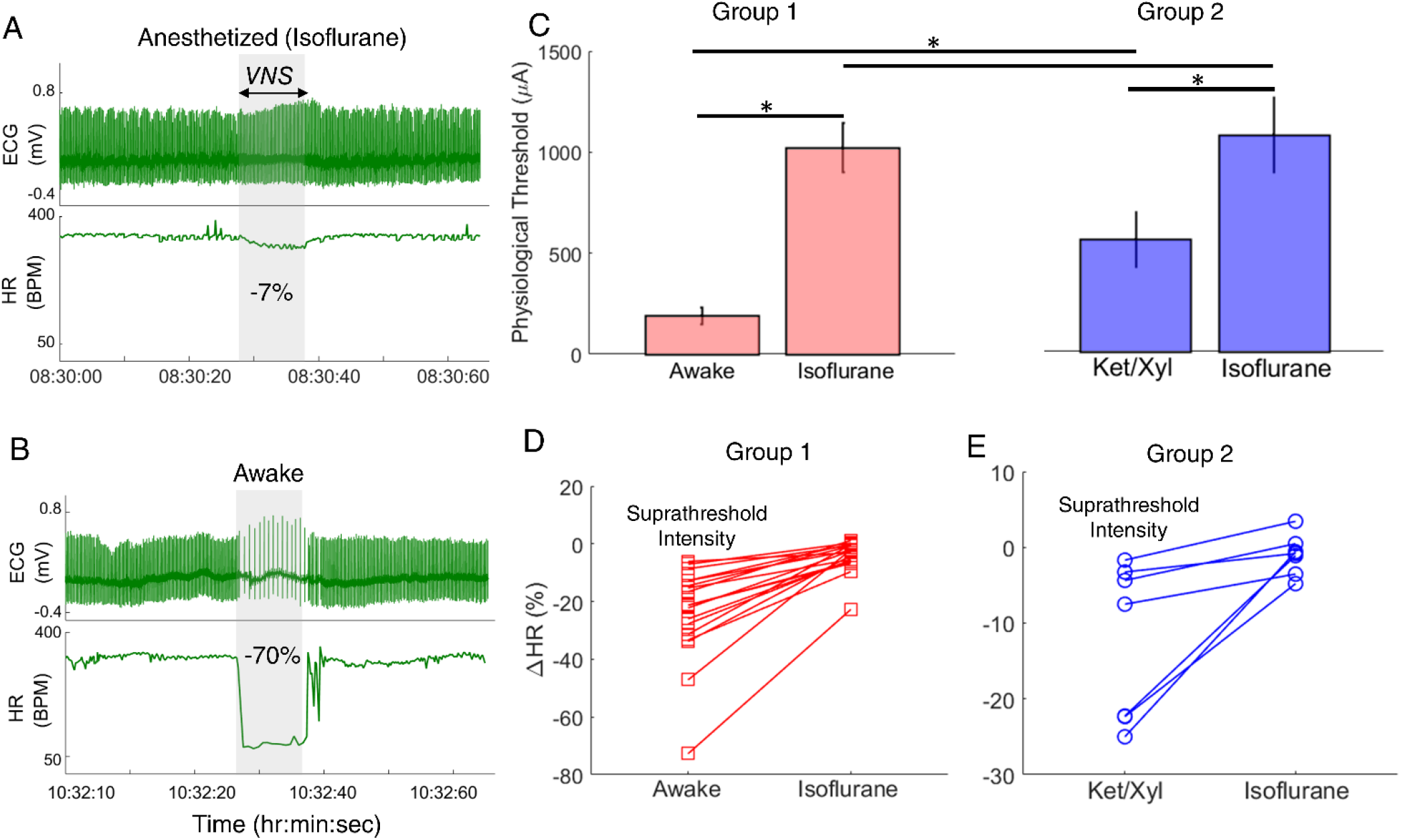
Effect of anesthesia on heart rate threshold of VNS. (A) Example in which VNS was delivered when the animal was anesthetized with isoflurane and stimulus-elicited change in heart rate (ΔHR) was registered. The grey shaded area represents the duration of stimulation. Parameters used were 1000uA, 100us, 30Hz. (B) The same animal as in (A) received the same stimulus after it was allowed to recover from isoflurane for at least 2 hours. (C) Left panel: Mean (±SEM) heart rate thresholds of VNS during wakefulness and during anesthesia with isoflurane (group 1: 23 experiments in 5 animals) (*p*<0.001, Paired *t*-test). Right panel: Mean (±SEM) heart rate thresholds of VNS during anesthesia with ketamine/xylazine and with isoflurane (group 2: 7 experiments in 5 animals) (*p*<0.01, Paired *t*-test). (D) Heart rate responses to VNS of the same suprathreshold intensities in each of the animals in group 1 (awake vs. isoflurance, left panel) and in group 2 (ketamine/xylazine vs. isoflurance, right panel).

## Discussion

In animal studies of vagus nerve stimulation (VNS), determining stimulation threshold is commonly one of the first steps in an experiment, as it defines the minimal effective “dose”. Expressing stimulation dose in units of threshold removes variability between experimental conditions or between animals (fig. 7). The selection of the type of threshold depends on the study outcome: “neural threshold” (NT), the minimal intensity that elicits nerve activity, is appropriate when nerve action potentials are studied^32^, whereas “physiological threshold” (PT), the minimal intensity that produces a physiological response, is more suitable when the physiological effects of VNS are of interest^33, 34^.

**Figure 7.**
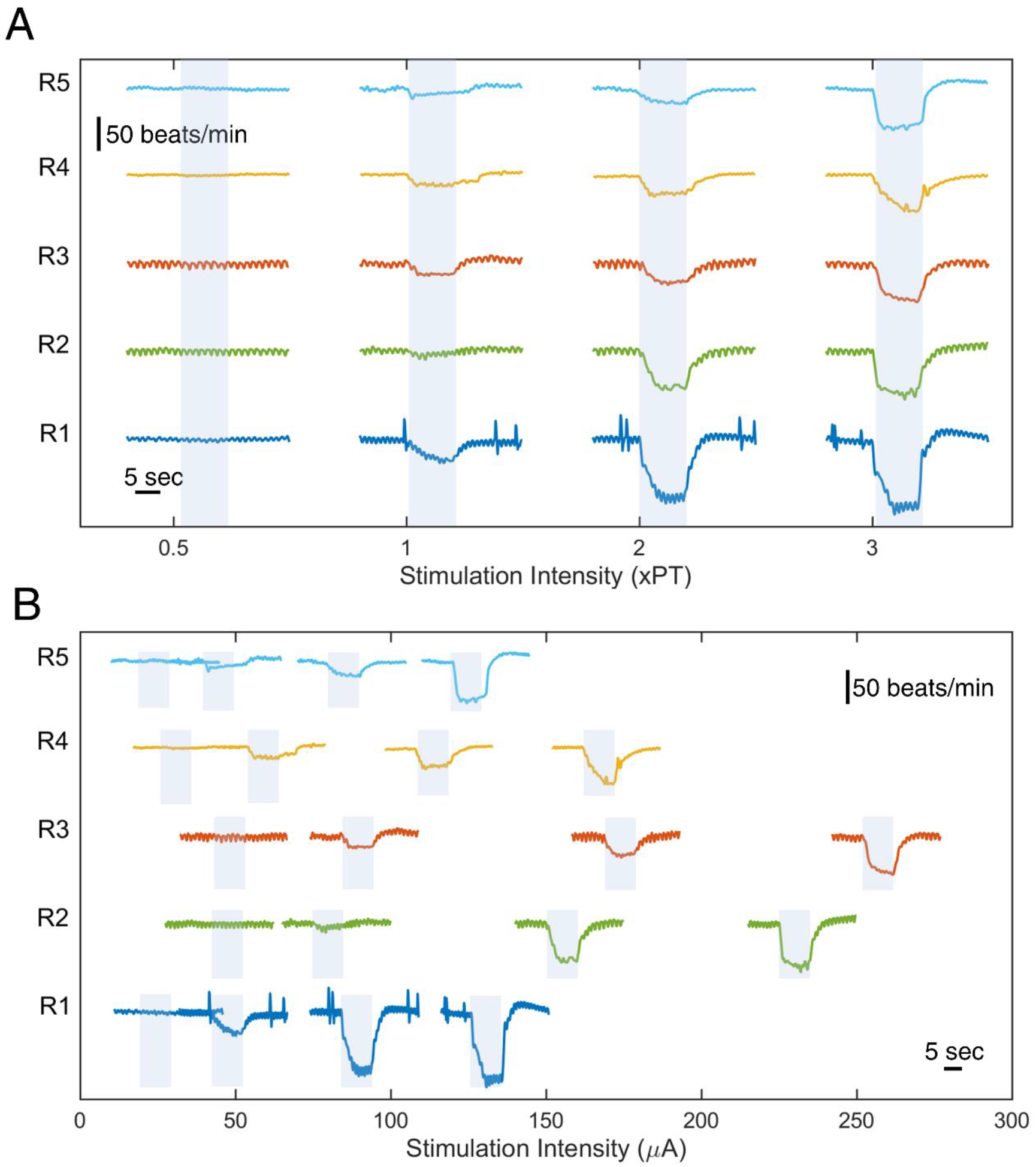
VNS-elicited changes in heart rate at different normalized or absolute stimulus intensities. (A) Change in heart rate at different normalized intensities (expressed in units of threshold intensity), in 5 animals. The colored shaded areas represent the duration of stimulation. (B) Change in heart rate at different absolute intensities, in the same 5 animals.

In our study, we determined both NT and several types of PT to cervical VNS in laboratory rats. PT was determined with respect to several different physiological responses, which followed a sequence as stimulus intensity was increased: increase in blood pressure (BP) and drop in breathing rate (BR) typically occur at lowest intensities, drop in heart rate (HR), and a resulting drop in BP (under isoflurane) or increase in BP (awake conditions), at intermediate intensities, and finally apnea, along with drop in HR and BP, at high intensities (table. III). Interestingly, the same sequence of physiological effects with increasing intensity is seen in both anesthetized and awake animals (fig. 4), something reported for the first time, to the best of our knowledge. This means that anesthesia does not alter the sequence of physiological effects of VNS as stimulus intensity increases.

This sequence of physiological effects can be explained by the stimulus intensity-dependent engagement of vagal fibers by VNS, which follows a size principle: larger fibers have relatively low thresholds and smaller fibers have high thresholds^35^. At low intensities, activation of larger A-delta fibers from lung stretch receptors may be responsible for the drop in breathing rate, as part of the Herring-Breuer reflex^36–38^. At those intensities, afferent A-delta fibers from aortic baroreceptors^39^ are also activated. Isolated activation of those fibers is known to elicit a drop in BP^40^, in contrast to the increase in BP we documented at low intensities. It is likely that in our case, activation of A-delta fibers happens concurrently with co-activation of efferent sympathetic fibers, of which there is a small number in the vagus^41^, resulting in vasoconstriction and increase in BP. It is also likely that engagement of afferent A-delta fibers initiates vagal-sympathetic reflexes resulting in an increase in the sympathetic tone to vessels, resulting in vasoconstriction, and to the heart, resulting in tachycardia^34^. Afferent fiber activation happening at these low intensities could also centrally suppress the parasympathetic tone to the heart^33, 34^. Such vagal-sympathetic reflexes are mediated centrally by direct connections from the vagal sensory nucleus of the solitary tract (NTS) to the dorsal motor nucleus (DMN) of the vagus and sympathetic ganglia^34, 42, 43^. As stimulus intensity increases further and cardio-inhibitory B-fibers become more engaged, their cardioinhibitory action at the heart dominates, and drops in HR and BP ensues^33, 34^ (fig. 4). Interestingly, in awake conditions, a similar drop in HR is observed but with an increase in SAP (table. III). This difference in response between anesthetized and awake conditions is possibly due to the fact that isoflurane suppresses sympathetic activity which resulted in the absence of an increase in SAP under isoflurane^44, 45^. Finally, activation of smaller C-fibers at highest stimulus intensities in rats produces apnea^38, 46, 47^.

On a single animal-basis, NT and PT are positively and highly correlated (fig. 3C); this finding may be used to estimate NT by measuring PT, under specific conditions. In general, measuring PT is much faster and easier than determining NT, which involves the step of recording stimulus-evoked compound action potentials (eCAPs). We found that NT ranged between 14-40 μA (single pulses of 100 μs pulse width) and PT between 32-150 μA (trains of 100 μs-long pulses at 30 Hz pulsing frequency). The higher values for PT than for NT are expected, given the additional anatomical elements involved in the determination of PT and have been described before^10, 35^. This difference can be explained by the neural components engaged in their measurement. For NT, those include the electrode-nerve interface and the level of excitability of the stimulated nerve fibers. Determining PT in addition, involves synaptic transmission at the ganglionic synapse, the level of excitability of the corresponding ganglionic cells, and the postganglionic synapse or circuit at the end-organ, as well as the state, at the time of stimulation, of the end-organ itself (fig. 8). It follows that, due to synaptic temporal summation, PT should be dependent on pulsing frequency. Indeed, we found that the physiological responses to VNS trains are different between 2 pulsing frequencies (1 Hz vs. 30 Hz), even though the eCAP responses to individual pulses of those same trains are largely similar (table I). We hypothesized that since a physiological response relies both on nerve activation and engagement of the neural circuit controlling the end organ (sinus node of the heart, baroreflex or respiratory center in the brain): (a) NT should be lower than PT, (b) PT, but not NT, should be dependent on pulsing frequency.

**Figure 8.**
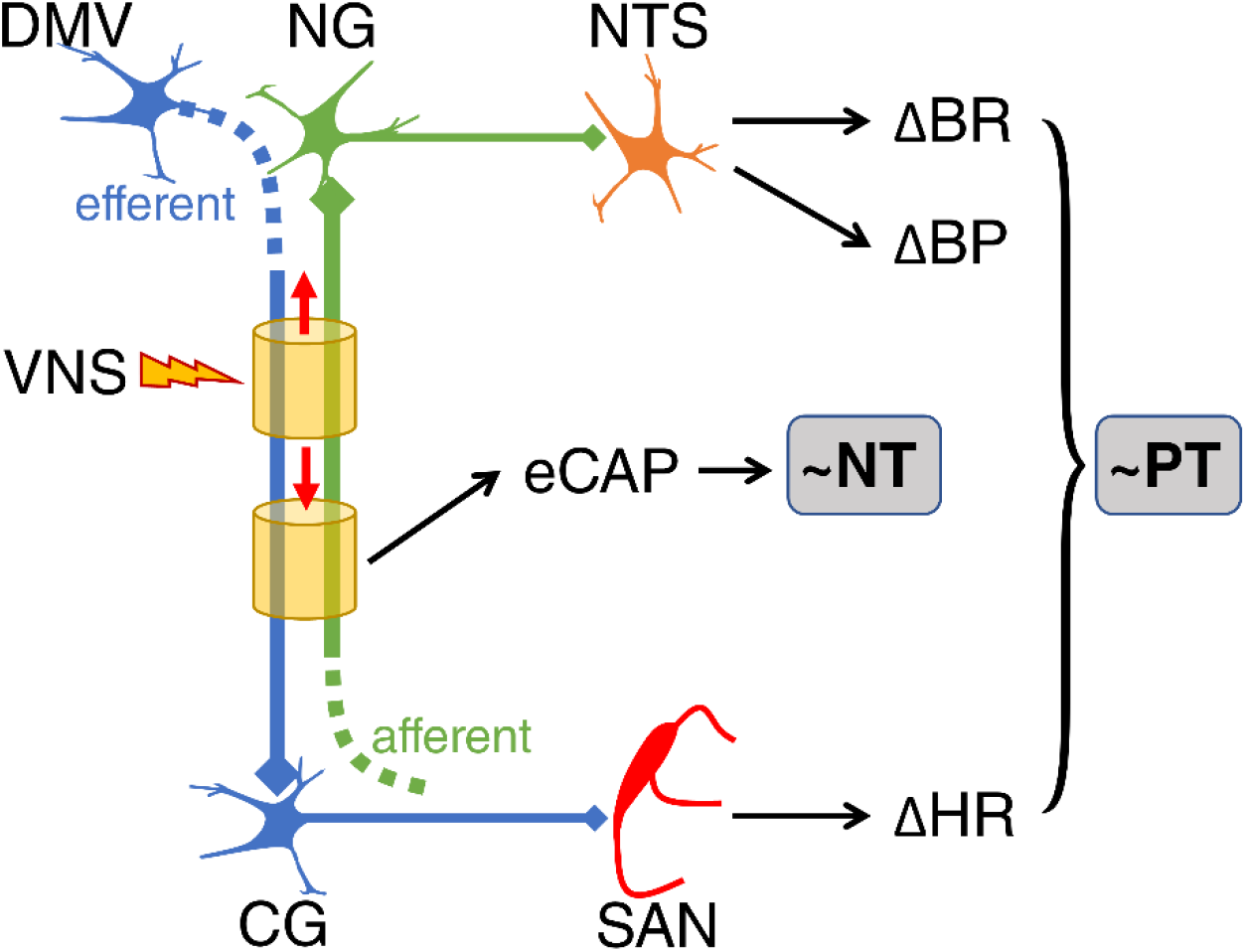
Circuit mechanisms involved in the determination of “neural threshold” (NT) and “physiological threshold” (PT) intensities of VNS. NT depends on the engagement of afferent and efferent vagal nerve fibers by stimuli, as measured by stimulus-evoked compound action potentials (eCAPs). PT depends on the engaged neural pathway and end-organ. Heart rate changes (ΔHR) occur when efferent axons of cells in the dorsal motor nucleus of the vagus (OMV) are engaged at a level at which they depolarize cells of cardiac ganglia (CG) that in turn project to the sino-atrial node (SAN). Breathing rate changes (ΔBR) and blood pressure changes (ΔBP) occur when afferent axons of pseudo-unipolar cells of the vagal ganglia (VG) are engaged at a level at which they depolarize cells in the nucleus of the solitary tract (NTS), engaging corresponding neural reflexes. Factors related to the interface between the stimulating electrode and the vagus nerve affects both NT and PT, whereas factors related to anesthesia affect primarily the PT, due to the involvement of synaptic transmission in the latter.

Adequate insulation of an electrode during in-vivo implantation is required to focus the resulting electrical field to the targeted nerve and to limit current spread and activation of non-targeted excitable tissues in the vicinity of the electrode^48–51^. In our experiments, PT with extra insulation, in the form of a silicone layer completely surrounding the electrode-nerve interface, is 10 times smaller than PT without that insulation (Suppl. table I). In one animal, “non-insulated” PT is reduced to a similar level as “insulated” PT after dabbing the fluid accumulation around the interface (Suppl. table I), indicating that the higher intensities needed with non-insulated interfaces are likely due to current shunting between the electrode contacts. In long-term implants, the insulating material must be biocompatible^52^, or else, tissue reaction may compromise the nerve and the electrode-nerve interface^53^, hence affecting the performance of the implant.

We found that PT increases with implant age (fig. 5C). Similarly, the intensities required to elicit a given suprathreshold physiological response are significantly higher at a later age compared to an earlier implant age (fig. 5A, 5B). This gradual rise in PT is due to a multifactorial process involving inflammatory and fibrotic response to the presence of the electrode^18, 19^, nerve injury with loss of nerve fibers^54^, and degradation of electrode contacts or leads^20^. Electrode impedance reflects the electrical characteristics of the electrode-nerve interface and has been suggested to be a measure of its effectiveness and integrity^55^. In our study, in-vivo electrode impedance increases with implant age (fig. 5D), in agreement with other reports^56, 57^. Both PT and impedance increase with implant age, having a weak but significant correlation on a single measurement basis (fig. 5E). When impedance is high (greater than 200 kΩ), PT is always indeterminable (fig. 5E), indicating a non-functional implant, but measurable impedance does not always guarantee a functional implant^57^ (fig. 5E). In acute experiments, neither *ex-vivo* nor *in-vivo* electrode impedance is correlated with PT (Suppl. fig. S4). That may be because PT is closely related to the spatial relationships between the electrode contacts and the different nerve fiber types that produce the physiological responses, especially at the relatively low threshold intensities used in our study. The anatomical heterogeneity of the fiber composition of the cervical vagus between animals^58^, the presence or absence of aortic depressor nerve in the carotid sheath along with the vagus^59^, the exact orientation of the electrode and the resulting electric field in relation to the nerve^22^, are all significant determinants of PT and their contribution is not captured by measuring *in vivo* or *ex vivo* electrode impedance.

The use of general anesthesia is typically a prerequisite for acute rodent experiments. General anesthetic agents provide analgesia and loss of consciousness, primarily by acting on the central nervous system (CNS). Actions on the autonomic nervous system (ANS) have also been described, including actions on ganglionic neurons^60^, synaptic transmission^61–63^, and higher anesthesia concentration can also block the axonal action potentials^64, 65^. Additionally, general anesthesia reduces the receptors’ sensitivity for neurotransmitters^66, 67^. In our study, we found that the negative chronotropic effect of VNS is maximum in awake state, followed by ketamine-xylazine, and isoflurane (fig. 6C-6E). These differences between awake and anesthetized states are likely because of the effects of general anesthesia on ANS as explained above. Our findings are in agreement with a previous study comparing isoflurane and awake states, which shows that at a given stimulation intensity, the chronotropic response of VNS was more pronounced in awake conditions than under isoflurane (figure 5 of Seagard et al^68^). We also found that PT is significantly lower in awake state compared to anesthesia (fig. 6C). The increase in PT under general anesthesia is also consistent with other peripheral nerve stimulation studies, which show the stimulation threshold of muscle contraction (end-organ) increases with general anesthesia, possibly as a result of inhibition of nicotinic acetylcholine receptors at the neuromuscular junction^69–71^. In addition to the effect of anesthesia on ANS, they also affect the cardiovascular system, specifically the heart. In particular, isoflurane has known to depress the cardiovascular system by inhibiting ion-channels on cardiac muscles and pacemaker. The channels it inhibits are the sodium channels responsible for generating action potentials^72^, the voltage-gated cardiac calcium channel responsible for generating pacemaker potentials^73^, and the voltage-gated potassium channels that delay the recovery from the action potential^74^. On the other hand, ketamine is an N-methyl-D-aspartate (NMDA) antagonist, which can also modulate parasympathetic cardiac activity by altering nicotinic cholinergic excitatory mechanisms^75^. The effects of ketamine on ANS, especially vagal tone, is controversial. One study indicated that ketamine does not affect the vagal tone, such as its effects on ganglion and synaptic potential transmission is limited^76^, while the other stated that it affects vagal tone by stimulating vagal fibers^77^. Additionally, xylazine is an alpha-2 adrenergic receptor antagonist that can cause central nervous system (CNS) depression^78^. We found out that physiological thresholds are significantly lower under ketamine-xylazine compare to isoflurane (fig. 6C); these findings can be explained by the fact that isoflurane has been shown to have a more pronounced effect on ANS and the heart compare to ketamine-xylazine. Hence, a pronounced chronotropic effect of VNS would be expected with ketamine-xylazine compare to isoflurane. Therefore, the discrepancies in physiological thresholds observed in awake and anesthetized conditions are probably due to a combination of anesthetic agent’s effects on the autonomic nervous system, as well as on the heart (end-organ).

## Acknowledgement

All authors declare no conflict of interest.

## Notes

**Sources of financial support** This work was supported in part by a research grant to SZ from United Therapeutics Corporation.

**Conflict of interest statement** All authors declare no conflict of interest.

### Competing Interest Statement

The authors have declared no competing interest.

## References

1. Rush AJ, George MS, Sackeim HA, et al. Vagus nerve stimulation (VNS) for treatment-resistant depressions: a multicenter study. Biol Psychiatry. Feb 15 2000;47(4):276–286.

2. Mehanna R, Machado AG, Connett JE, Alsaloum F, Cooper SE. lntraoperative Microstimulation Predicts Outcome of Postoperative Macrostimulation in Subthalamic Nucleus Deep Brain Stimulation for Parkinson’s Disease. Neuromodulation. Jul 2017;20(5):456–463.

3. Shahwan A, Bailey C, Maxiner W, Harvey AS. Vagus nerve stimulation for refractory epilepsy in children: More to VNS than seizure frequency reduction. Epilepsia. 2009/05/01 2009;50(5):1220–1228.

4. Deuschl G, Schade-Brittinger C, Krack P, et al. A randomized trial of deep-brain stimulation for Parkinson’s disease. N Engl J Med. Aug 31 2006;355(9):896–908.

5. Kaniusas E, Kampusch S, Tittgemeyer M, et al. Current Directions in the Auricular Vagus Nerve Stimulation II - An Engineering Perspective. Front Neurosci. 2019;13:772.

6. Liu A, Voroslakos M, Kronberg G, et al. Immediate neurophysiological effects of transcranial electrical stimulation. Nat Commun. Nov 30 2018;9(1):5092.

7. Berenyi A, Belluscio M, Mao D, Buzsaki G. Closed-loop control of epilepsy by transcranial electrical stimulation. Science. Aug 10 2012;337(6095):735–737.

8. Bucksot JE, Wells AJ, Rahebi KC, et al. Flat electrode contacts for vagus nerve stimulation. PLoS One. 2019;14(11):e0215191.

9. Krames ES, Hunter Peckham P, Rezai A, Aboelsaad F. Chapter 1 - What Is Neuromodulation? In: Krames ES, Peckham PH, Rezai AR, eds. Neuromodulation. San Diego: Academic Press; 2009:3–8.

10. Yoo PB, Liu H, Hincapie JG, Ruble SB, Hamann JJ, Grill WM. Modulation of heart rate by temporally patterned vagus nerve stimulation in the anesthetized dog. Physiol Rep. Feb 2016;4(2).

11. Bally JF, Vargas Ml, Horvath J, et al. Localization of Deep Brain Stimulation Contacts Using Corticospinal/Corticobulbar Tracts Stimulation. Front Neural. 2017;8:239.

12. Mekhail N, Levy RM, Deer TR, et al. Long-term safety and efficacy of closed-loop spinal cord stimulation to treat chronic back and leg pain (Evoke): a double-blind, randomised, controlled trial. Lancet Neural. Feb 2020;19(2):123–134.

13. Sabbah HN. Electrical vagus nerve stimulation for the treatment of chronic heart failure. Cleve Clin J Med. Aug 2011;78 Suppl 1:S24–29.

14. Legatt AD. Motor Evoked Potentials. In: Aminoff MJ, Daroff RB, eds. Encyclopedia of the Neurological Sciences (Second Edition). Oxford: Academic Press; 2014:111–114.

15. Campbell CM, Jamison RN, Edwards RR. Psychological screening/phenotyping as predictors for spinal cord stimulation. Curr Pain Headache Rep. Jan 2013;17(1):307.

16. Seno SI, Shimazu H, Kogure E, Watanabe A, Kobayashi H. Factors Affecting and Adjustments for Sex Differences in Current Perception Threshold With Transcutaneous Electrical Stimulation in Healthy Subjects. Neuromodulation. Jul 2019;22(5):573–579.

17. Ahmed U, Chang YC, Cracchiolo M, et al. Anodal block permits directional vagus nerve stimulation. Sci Rep. Jun 8 2020;10(1):9221.

18. Moazzam Z, Paquette J, Duke AR, Khodaparast N, Yoo PB. Feasibility of Long-term Tibial Nerve Stimulation Using a Multi-contact and Wirelessly Powered Neurostimulation System Implanted in Rats. Urology. 2017/04/01/ 2017;102:61–67.

19. Coleman DL, King RN, Andrade JD. The foreign body reaction: a chronic inflammatory response. J Biomed Mater Res. Sep 1974;8(5):199–211.

20. Caldwell R, Street MG, Sharma R, Takmakov P, Baker B, Rieth L. Characterization of Parylene-C degradation mechanisms: In vitro reactive accelerated aging model compared to multiyear in vivo implantation. Biomaterials. Feb 2020;232:119731.

21. Howell B, Naik S, Grill WM. Influences of interpolation error, electrode geometry, and the electrode-tissue interface on models of electric fields produced by deep brain stimulation. IEEE Trans Biomed Eng. Feb 2014;61(2):297–307.

22. Aristovich K, Donega M, Fjordbakk C, et al. Model-based geometrical optimisation and in vivo validation of a spatially selective multielectrode cuff array for vagus nerve neuromodulation. arXiv preprint arXiv:1903.12459. 2019.

23. Zander H, Graham R, Anaya CJ, Lempka S. Anatomical and technical factors affecting the neural response to epidural spinal cord stimulation. J Neural Eng. May 4 2020.

24. Tsui BC. The effects of general anaesthesia on nerve-motor response characteristics (rheobase and chronaxie) to peripheral nerve stimulation. Anaesthesia. 2014/04/01 2014;69(4):374–379.

25. Handforth A, DeGiorgio CM, Schachter SC, et al. Vagus nerve stimulation therapy for partial-onset seizures: a randomized active-control trial. Neurology. Jul 1998;51(1):48–55.

26. Arie JE, Carlson KW, Mei L. Investigation of mechanisms of vagus nerve stimulation for seizure using finite element modeling. Epilepsy Research. 2016/10/01/ 2016;126:109–118.

27. De Ferrari GM, Tuinenburg AE, Ruble S, et al. Rationale and study design of the NEuroCardiac TherApy foR Heart Failure Study: NECTAR-HF. Eur J Heart Fail. Jun 2014;16(6):692–699.

28. Aranow C, Lesser M, Mackay M, et al. Engaging the Cholinergic Anti-Inflammatory Pathway By Stimulating the Vagus Nerve Reduces Pain and Fatigue in Patients with SLE. Paper presented at: ARTHRITIS & RHEUMATOLOGY2018.

29. Koopman FA, Chavan SS, Miljko S, et al. Vagus nerve stimulation inhibits cytokine production and attenuates disease severity in rheumatoid arthritis. Proc Natl Acad Sci U S A. Jul 19 2016;113(29):8284–8289.

30. Masi EB, Valdes-Ferrer SI, Steinberg BE. The vagus neurometabolic interface and clinical disease. Int J Obes (Land). Jun 2018;42(6):1101–1111.

31. Gribi S, du Bois de Dunilac S, Ghezzi D, Lacour SP. A microfabricated nerve-on-a-chip platform for rapid assessment of neural conduction in explanted peripheral nerve fibers. Nat Commun. Oct 23 2018;9(1):4403.

32. Vuckovic A, Tosato M, Struijk JJ. A comparative study of three techniques for diameter selective fiber activation in the vagal nerve: anodal block, depolarizing prepulses and slowly rising pulses. J Neural Eng. Sep 2008;5(3):275–286.

33. Ardell JL, Nier H, Hammer M, et al. Defining the neural fulcrum for chronic vagus nerve stimulation: implications for integrated cardiac control. J Physiol. Nov 15 2017;595(22):6887–6903.

34. Ardell JL, Rajendran PS, Nier HA, KenKnight BH, Armour JA. Central-peripheral neural network interactions evoked by vagus nerve stimulation: functional consequences on control of cardiac function. Am J Physiol Heart Gire Physiol. Nov 15 2015;309(10):H1740–1752.

35. McAllen RM, Shafton AD, Bratton BO, Trevaks D, Furness JB. Calibration of thresholds for functional engagement of vagal A, Band C fiber groups in vivo. Bioelectron Med (Land). Jan 2018;1(1):21–27.

36. Hayashi F, Coles SK, McCrimmon DR. Respiratory neurons mediating the Breuer-Hering reflex prolongation of expiration in rat. J Neurosci. Oct 15 1996;16(20):6526–6536.

37. Paintal AS. Vagal sensory receptors and their reflex effects. Physiol Rev. Jan 1973;53(1):159–227.

38. Chang YC, Cracchiolo M, Ahmed U, et al. Quantitative estimation of nerve fiber engagement by vagus nerve stimulation using physiological markers. Brain Stimul. Sep 18 2020;13(6):1617–1630.

39. Fan W, Andresen MC. Differential frequency-dependent reflex integration of myelinated and nonmyelinated rat aortic baroreceptors. Am J Physiol. Aug 1998;275(2):H632–640.

40. De Paula PM, Castania JA, Bonagamba LG, Salgado HC, Machado BH. Hemodynamic responses to electrical stimulation of the aortic depressor nerve in awake rats. Am J Physiol. Jul 1999;277(1):R31–38.

41. Seki A, Green HR, Lee TD, et al. Sympathetic nerve fibers in human cervical and thoracic vagus nerves. Heart Rhythm. Aug 2014;11(8):1411–1417.

42. Andresen MC, Kunze DL, Mendelowitz D, Armour JA, Ardell JL. Basic and Clinical Neurocardiology. 2004.

43. Sawchenko PE. Central connections of the sensory and motor nuclei of the vagus nerve. J Auton Nerv Syst. Oct 1983;9(1):13–26.

44. Wood M. Effect of general anesthesia on modulation of sympathetic nervous system function. Adv Pharmacol. 1994;31:449–458.

45. Skovsted P, Sapthavichaikul S. The effects of isoflurane on arterial pressure, pulse rate, autonomic nervous activity, and barostatic reflexes. Can Anaesth Soc J. May 1977;24(3):304–314.

46. Carr MJ, Undem BJ. Bronchopulmonary afferent nerves. Respirology. Sep 2003;8(3):291–301.

47. Bozler E, Burch BH. Role of the vagus in the control of respiration. Am J Physiol. Aug 1951;166(2):255–261.

48. Fan Y, Jiang E, Hahka T, Chen QH, Yan J, Shan Z. Orexin A increases sympathetic nerve activity through promoting expression of proinflammatory cytokines in Sprague Dawley rats. Acta Physiol (Oxf). Feb 2018;222(2).

49. Lasztoczi B, Klausberger T. Distinct gamma oscillations in the distal dendritic fields of the dentate gyrus and the CA1 area of mouse hippocampus. Brain Struct Funct. Sep 2017;222(7):3355–3365.

50. Kawada T, Turner MJ, Shimizu S, Fukumitsu M, Kamiya A, Sugimachi M. Aortic depressor nerve stimulation does not impede the dynamic characteristics of the carotid sinus baroreflex in normotensive or spontaneously hypertensive rats. Am J Physiol Regul lntegr Comp Physiol. May 1 2017;312(5):R787–R796.

51. Nicolai EN, Settell ML, Knudsen BE, et al. Sources of off-target effects of vagus nerve stimulation using the helical clinical lead in domestic pigs. J Neural Eng. Jul 24 2020;17(4):046017.

52. Szostak KM, Grand L, Constandinou TG. Neural Interfaces for lntracortical Recording: Requirements, Fabrication Methods, and Characteristics. Front Neurosci. 2017;11:665.

53. Anderson JM, Rodriguez A, Chang OT. Foreign body reaction to biomaterials. Semin lmmunol. Apr 2008;20(2):86–100.

54. Grill WM, Mortimer JT. Neural and connective tissue response to long-term implantation of multiple contact nerve cuff electrodes. J Biomed Mater Res. May 2000;50(2):215–226.

55. Kappenman ES, Luck SJ. The effects of electrode impedance on data quality and statistical significance in ERP recordings. Psychophysiology. Sep 2010;47(5):888–904.

56. Yaghouby F, Shafer B, Vasudevan S. A rodent model for long-term vagus nerve stimulation experiments. Bioelectronics in Medicine. 2019/06/01 2019;2(2):73–88.

57. Mughrabi IT, Hickman J, Jayaprakash N, et al. An implant for long-term cervical vagus nerve stimulation in mice. bioRxiv. 2020:2020.2006.2020.160473.

58. Settell ML, Pelot NA, Knudsen BE, et al. Functional vagotopy in the cervical vagus nerve of the domestic pig: implications for the study of vagus nerve stimulation. J Neural Eng. Apr 9 2020;17(2):026022.

59. Krieger EM, Marseillan RF. Aortic Depressor Fibers in the Rat: An Electrophysiological Study. Am J Physiol. Oct 1963;205:771–774.

60. Boban N, McCallum JB, Schedewie HK, Boban M, Kampine JP, Bosnjak ZJ. Direct comparative effects of isoflurane and desflurane on sympathetic ganglionic transmission. Anesth Analg. Jan 1995;80(1):127–134.

61. Maciver MB. Anesthetic agent-specific effects on synaptic inhibition. Anesth Analg. Sep 2014;119(3):558–569.

62. Matthews EK, Quilliam JP. Effects of Central Depressant Drugs Upon Acetylcholine Release. Br J Pharmacol Chemother. Apr 1964;22:415–440.

63. Pocock G, Richards CD. The action of volatile anaesthetics on stimulus-secretion coupling in bovine adrenal chromaffin cells. Br J Pharmacol. Sep 1988;95(1):209–217.

64. Seeman P. The membrane actions of anesthetics and tranquilizers. Pharmacol Rev. Dec 1972;24(4):583–655.

65. Mikulec AA, Pittson S, Amagasu SM, Monroe FA, Maciver MB. Halothane depresses action potential conduction in hippocampal axons. Brain Res. Jun 15 1998;796(1-2):231–238.

66. Richards CD, Smaje JC. Anaesthetics depress the sensitivity of cortical neurones to L-glutamate. Br J Pharmacol. Nov 1976;58(3):347–357.

67. Sawada S, Yamamoto C. Blocking action of pentobarbital on receptors for excitatory amino acids in the guinea pig hippocampus. Exp Brain Res. 1985;59(2):226–231.

68. Seagard JL, Elegbe EO, Hopp FA, et al. Effects of isoflurane on the baroreceptor reflex. Anesthesiology. Dec 1983;59(6):511–520.

69. Paul M, Fokt RM, Kindler CH, Dipp NC, Yost CS. Characterization of the interactions between volatile anesthetics and neuromuscular blockers at the muscle nicotinic acetylcholine receptor. Anesth Analg. Aug 2002;95(2):362–367, table of contents.

70. Vanlinthout LE, Booij LH, van Egmond J, Robertson EN. Effect of isoflurane and sevoflurane on the magnitude and time course of neuromuscular block produced by vecuronium, pancuronium and atracurium. Br J Anaesth. Mar 1996;76(3):389–395.

71. Tsui BC. The effects of general anaesthesia on nerve-motor response characteristics (rheobase and chronaxie) to peripheral nerve stimulation. Anaesthesia. Apr 2014;69(4):374–379.

72. Rehberg B, Xiao YH, Duch OS. Central nervous system sodium channels are significantly suppressed at clinical concentrations of volatile anesthetics. Anesthesiology. May 1996;84(5):1223–1233; discussion 1227A.

73. Bosnjak ZJ, Aggarwal A, Turner LA, Kampine JM, Kampine JP. Differential effects of halothane, enflurane, and isoflurane on Ca2+ transients and papillary muscle tension in guinea pigs. Anesthesiology. Jan 1992;76(1):123–131.

74. Huneke R, Jungling E, Skasa M, Rossaint R, Luckhoff A. Effects of the anesthetic gases xenon, halothane, and isoflurane on calcium and potassium currents in human atrial cardiomyocytes. Anesthesiology. Oct 2001;95(4):999–1006.

75. lrnaten M, Wang J, Venkatesan P, et al. Ketamine inhibits presynaptic and postsynaptic nicotinic excitation of identified cardiac parasympathetic neurons in nucleus ambiguus. Anesthesiology. Mar 2002;96(3):667–674.

76. Yamamura T, Kimura T, Furukawa K. Effects of halothane, thiamylal, and ketamine on central sympathetic and vagal tone. Anesth Analg. Feb 1983;62(2):129–134.

77. Inoue K, Arndt JO. Efferent vagal discharge and heart rate in response to methohexitone, althesin, ketamine and etomidate in cats. Br J Anaesth. Oct 1982;54(10):1105–1116.

78. Flecknell P. Laboratory Animal Anaesthesia. Fourth ed: Elsevier; 2015.

