## Supplemental File # 1 for "Implant- and anesthesia-related factors affecting threshold intensities for vagus nerve stimulation"

### Methods

#### Electrode Fabrication

##### Flex Electrode Fabrication

The electrodes used for this study were designed for acute and chronic implantation, and made using microfabrication processes. The *Flex* electrodes are comprised of spin-cast polyimide foils with a sputter deposited bond-pad and trace metallization, and sputter deposited iridium oxide (IrOx) electrode sites. These IrOx electrode sites enable low electrode impedances while maintaining stable stimulation characteristics. The electrode sites have an area of 1,418×167 $\mu m^2$, and center-to-center spacing between the electrodes of 1 mm. The electrode fabrication process involves spin coating a 4” *p*-type silicon wafers with polyimide (PI) resin (HD Microsystems PI 2611, Parlin, NJ), which is cured at 300 °C in $N_2$. This generates a ~7 µm thick bottom dielectric. Liftoff photoresist (AZ 9260, Microchemicals, Ulm, Germany) is then spun onto the wafers and patterned using typical contact photolithographic processing, which defines the metallization for the bond pads, traces, and the base layer for the electrode sites. The pad and trace metallization layers are then sputter deposited using conditions optimized to minimize residual stress, generate stable films, and have good adhesion to the polyimide substrate. This metallization structure is Ti/Pt/Au, which is used to enable multiple methods to connect to the devices including soldering, conductive adhesive, and wire bonding. The liftoff resist is then stripped using acetone, followed by an IPA rinse, and cleaning with spin-rinse-dryer (SRD). A second liftoff photoresist to pattern the IrOx for the electrode sites is spin-coated onto the wafers using identical procedure. The IrOx stack is comprised of Ti/Ir/IrOx layers to achieve good adhesion and film stability. The resist is stripped using acetone, IPA, and SRD processes, which is similar to the prior process. A layer of PI for the top dielectric with identical characteristics to the previous PI layer is then spun on the wafer, and cured at 300 °C in $N_2$. The PI is then patterned to 1) open the electrode sites, 2) open the bond pads, and 3) define the device outline. Two layers of photoresist are spun and baked on the wafer yielding a resist layer ~20 μm thick, to sufficiently protect the polyimide during etching. The devices are then etched using an RIE process (Plasmalab 100, Oxford Instruments, Bristol, UK) using $O_2$ process gas at 10 mTorr using 300 W. The trace and electrode metallizations act as etch stop layers for those openings, while the process also etches through both top and bottom dielectric layers of PI to define/isolate the shape of the electrodes. The electrodes are mechanically released from the wafer using tweezers or a scalpel.

#### Acute Electrode Integration

Electrodes for acute animal preparations are integrated with leads and encapsulated. Lead wires are silicone-insulated stranded silver-plated copper wires (AS155-36, Cooner Wire, Chatsworth, CA), which are mechanically stripped. The wires are soldered onto the released flexible electrodes using lead-free solder with water-soluble flux (Indium Corp., Clinton, NY). The electrodes and leads are thoroughly rinsed in DI water and dried. A small amount of medical grade epoxy (Loctite M-31CL, Rockhill, CT) is used to mechanically stabilize and strengthen the lead attachment. The area around the lead attachment region is then encapsulated with medical grade silicone (NuSil MED 4211, Carpenteria, CA) as the final encapsulation. The impedances of the devices are measured in phosphate buffered saline (PBS) at room temperature using a 3-electrode configuration. A Pt wire counter electrode, and Ag|AgCl reference electrode, are used with a Gamry 600+ EIS system (Gamry Instruments, Warminster, PA). Typical electrode impedances in saline range from 0.5 to 1 kΩ at the frequency of 1 kHz. Photographs of completed bipolar and tripolar electrodes are presented in Fig. XXX.

#### Chronic Electrode Integration

Electrodes for chronic animal preparation use more robust leads and connector systems to facilitate long-term in-dwelling use. The leads use a helically wrapped wire approach similar to what is commonly used for medical devices, and based on experiences with Utah arrays in peripheral nerves. The connector system involves using an Omnetics Nanostrip connector on a printed circuit board (PCB), and a 3D printed Connector Protector to protect the junction of leads to the PCB.

The first aspect of the integration approach in fabrication of the lead. A set of three Pt-Ir wires using 9-strand wires with PTFE (Teflon) insulation that are 100 μm in outer diameter (Medwire, Mt. Vernon, NY) are helically wrapped around a 350 μm diameter rod to the appropriate length for the 5 cm lead body. A custom 2-part mold with a channel diameter of 1.0 mm is pre-loaded with NuSil MED-4211, and the helical wires a placed into the mold, which is then clamped together. The silicone is cured in an oven at 60 °C for 45 minutes. Two of the lead wires are attached to a flex electrode, and two of the wires are used as ground and/or reference wires for stimulation and recording. The ground and reference wires are deinsulated ~5 mm to form a low impedance (~200 Ω) connection to physiological fluids.

A 2-channel flex electrode with ~800 Ω electrode impedances is attached to the distal end of the lead with a combination of soldering and encapsulation. A small amount of lead-free SAC 305 alloy solder paste with water-soluble flux (Chip-Quik, Ancaster, ON) is dispensed on the bond pads, and the connection is made by placing the appropriate wire from the helical lead preform on the pad followed by using a fine-tip soldering iron to make the connection. After the two wires are connected, the electrode and lead are thoroughly rinsed in DI water to clean and remove the solder flux and any residual solder paste. A small amount of Loctite M-31CL (Rockhill, CT) medical-grade epoxy cured at 60 °C for >45 minutes is then used to mechanically stabilize the junction between wires and flex electrodes. The bond pad region of the flex electrodes is then overmolded in NuSil MED-4211 for optimized mechanical and encapsulation properties. This process is repeated to make 2 electrodes with leads.

After integration of the electrodes and leads, the electrodes are thermoformed such that they can wrap around the vagus nerve of the rat models. The electrode sites of a Flex electrode are wrapped around a 350 μm diameter tungsten rod as a mandrel for shaping the electrode. The Flex electrode is then heated with hot air from a Weller WTHA1N hot-air rework station (Besigheim, Germany) using 260 °C air with an airflow setting of 2. The hot air is applied 2 angles for 3 minutes at each angle.

The last steps of the integration process are to attach 2 electrode assemblies to an Omnetics connector assembly and attach a suture mesh used to help anchor the leads. An Omnetics A79018 nanostrip connector was integrated with a small custom PCB with a solder reflow process using SAC 305 solder with water-soluble flux, followed by a careful cleaning process in DI water. The base of the Omnetics connector was then mechanically stabilized and encapsulated on the PCB using Loctite M-31CL. The 2 electrode assemblies were then attached to the PCB using the same solder and cleaning processes as previously described, followed by applying a small quantity of epoxy to mechanically stabilize the joint. A Connector Protector made from 3D printed biocompatible Dental LT Clear Resin (Formlabs, Sommerville, MA) is then attached to the PCB to cover the attached wires on the PCB. The Connector Protector is used to generate a stiff and dimensionally stable surface to facilitate use of dental acrylic to attach the head-stage to the rat models skull, and mechanically protect the junction of the leads to the connector PCB. The Connector Protector is then back-filled with NuSil MED-4211 to further encapsulate the junction of the leads with the PCB, while maintaining consistent elastomeric mechanical properties with the helical lead. Lastly, surgical mesh suture (SurgicalMesh, Brookfield, CT) wings are attached to the distal aspect of the lead with NuSil 4211, which are used to anchor the electrodes near the nerve.

### Results

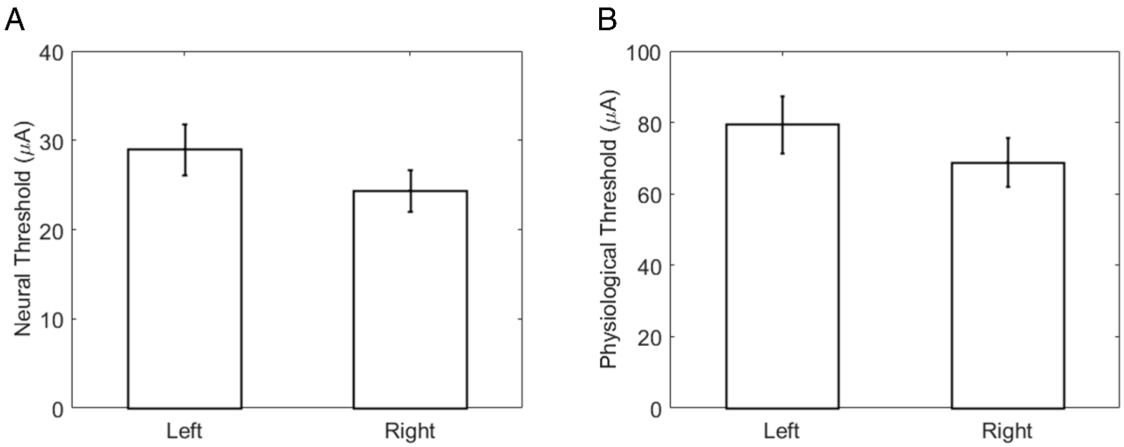


**Supplementary Figure S1. Threshold comparison of left and right vagus nerve stimulation in rats.**

A. Mean neural thresholds were compared for the right and left cervical VNS (n=8 for left, n=11 for right). Error bars represents SEM (*p* NS, Paired t-test).

B. Same as (A) for mean physiological thresholds (*p* NS, Paired t-test).

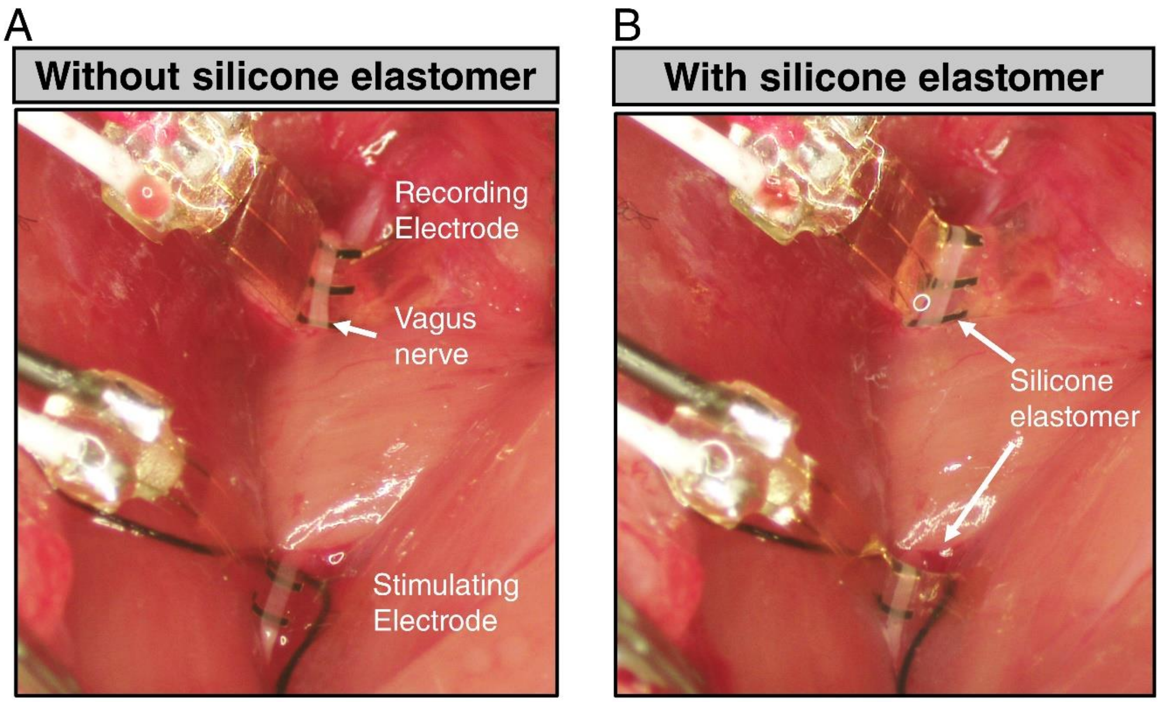


**Supplementary Figure S2. Neural interface with and without insulation.**

A. Acute implantation of two Flex electrodes on the rostral and caudal end of a right cervical vagus nerve in a rat. Image of a neural interface without silicone elastomer insulation.

B. Image of a neural interface with silicone elastomer insulation.

| Rat ID | Electrode | Physiological Thresholds (μA) | |
|---|---|---|---|
| | | Without silicone elastomer | With silicone elastomer |
| R1 | Cortec | 850 | 20 |
| R2 | Cortec | 1000 | 80 |
| R3 | Flex | 750 | 30 |
| R4 | Flex | 1000 | 110 |
| R5 | Flex | 500 | 30 |
| R6 | Flex | 750 | 50 |
| R7 | Flex | 80* | 60 |

**Supplementary Table 1. Effect of electrode insulation on the physiological threshold (PT) of VNS.** PT was first identified without silicone elastomer. In the same animal, PT was identified again after application of silicone elastomer. Asterisk indicates dabbing of excess fluid before stimulation.

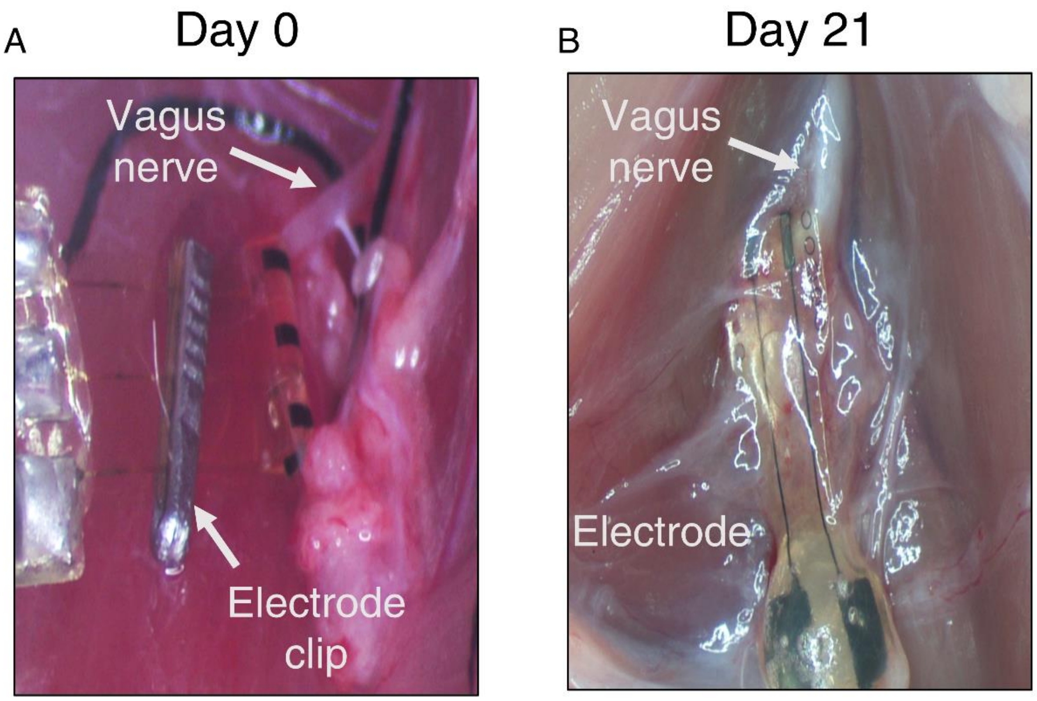


**Supplementary Figure S3. Gross view of cervical vagus nerve in rats.**

A. Gross image of implanted electrode around the left cervical vagus nerve on the day of surgery.

B. Image of the nerve taken during autopsy after 21 days of implantation. The electrode clip was removed to better visualize the foreign body response.

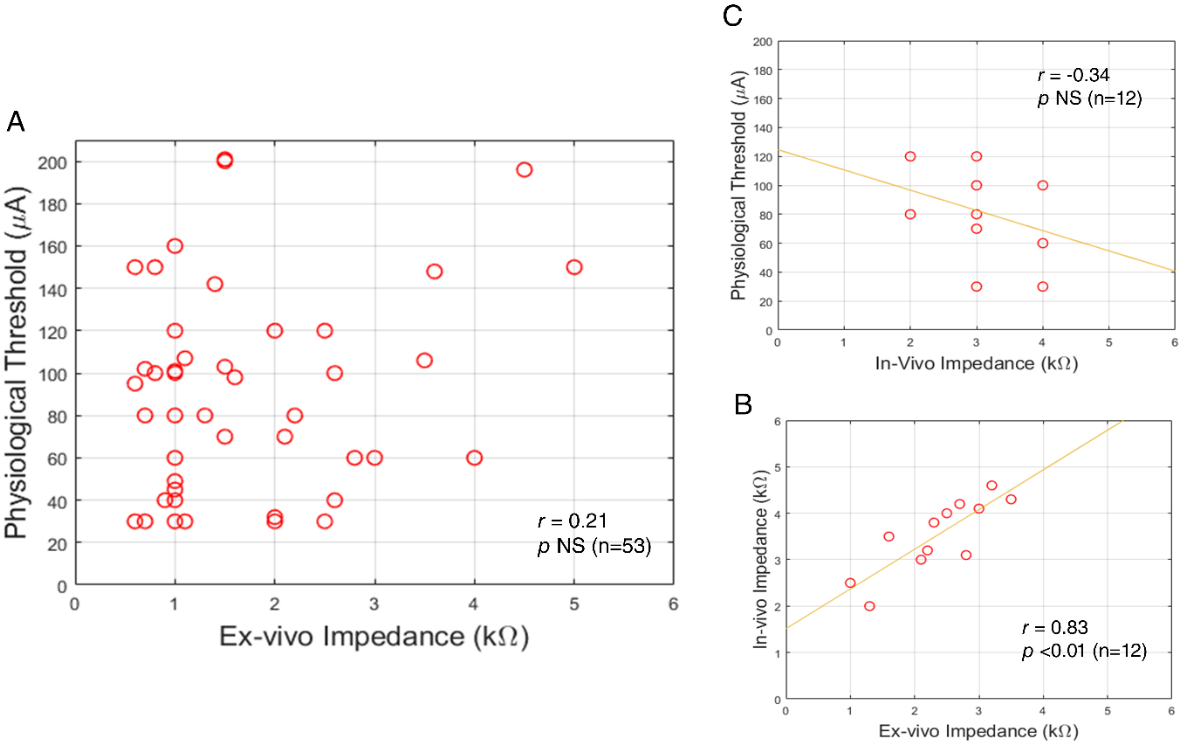


**Supplementary Figure S4. Correlation between physiological threshold and electrode impedance in acute VNS experiments.**

A. The scatterplot shows ex-vivo electrode impedance values with corresponding physiological thresholds in acute experiments. Each circle represents an experiment (total of 53 experiments from 53 animals; Pearson correlation coefficient: *r* = 0.21, *p* NS).

B. Scatterplot shows in-vivo electrode impedance values with corresponding physiological thresholds in acute experiments. Each circle represents an experiment (total of 12 experiments from 12 animals; Pearson correlation coefficient: *r* = 0.34, *p* NS).

C. Scatterplot showing ex-vivo and in-vivo impedances of individual electrodes. A linear line was best fitted based on the data plotted (total of 12 experiments from 12 animals; *r* = 0.83, *p* <0.001).
